# A holocentric pangenome links karyotype evolution to meiotic recombination

**DOI:** 10.64898/2026.01.17.700048

**Authors:** Meng Zhang, Stefan Steckenborn, Marco Castellani, Laia Marín-Gual, Amanda Souza Câmara, Laura A. Robledillo, Letícia Maria-Parteka, Nafiseh Sargheini, Magdalena Marek, Eduardo Chacón-Madrigal, Andrés Gatica Arias, Leonardo P. Félix, William W. Thomas, Bruno Huettel, Andrea Pedrosa-Harand, John T. Lovell, André L. L. Vanzela, Aurora Ruiz-Herrera, André Marques

## Abstract

Chromosomal fissions, fusions and whole-genome duplications propel genome evolution, yet their impact on meiotic recombination is obscured by the centromere constraint, since in monocentric species most large rearrangements are lethal^1–4^. Holocentric organisms, which distribute kinetochore activity along the entire chromosome, overcome this barrier and therefore offer a unique window onto the interplay between karyotype change and crossover control^2^. We assembled chromosome-scale genomes for 20 holocentric *Rhynchospora* species (including 56 haplotypes), representing all major clades of the genus, featuring satellite-based holocentromeres^5,6^, and integrated single-gamete crossover maps, high-resolution meiotic synapsis immunocytochemistry and Hi-C chromatin architecture. Breakpoint analysis shows that holocentromeric *Tyba* satellite arrays^6,7^ are recurrent hotspots for both chromosome fusions and fissions, contributing to the genus’s extraordinary chromosome number variation from 2*n* = 4 to 36. Crossover landscapes group into two apparent modes: strongly distal-biased versus irregularly distributed, which is correlated with divergent patterns of synapsis elongation. Moreover, crossover number scales with chromosome count and meiotic axis length. In contrast, crossover density per megabase is inversely related to chromosome length and to chromatin-loop size. We propose that chromosome fissions create karyotypes with smaller chromosomes folded into shorter loops, thereby increasing the axial substrate accessible for double-strand break formation and elevating recombination frequency. Together, our results provide a structural link between large-scale structural chromosome evolution and meiotic recombination through coupled changes in chromosome number, size, loop geometry, and synapsis dynamics.

## Introduction

Chromosomal rearrangements, including end-to-end reciprocal chromosome translocations (hereafter called ‘fusions’), fissions, and whole-genome duplications, are major drivers of genome evolution, species divergence, and adaptation^2–4,8,9^. These structural changes alter karyotypes and impact gene flow, recombination, and reproductive isolation^2,9,10^. While such changes are widespread, their consequences for chromosome behaviour and genetic diversity vary dramatically across taxa and chromosome architectures^2,9^.

In monocentric species, where chromosome segregation relies on a single centromere, large-scale rearrangements often compromise chromosome stability, leading to missegregation or even lethality^1,2^. In contrast, holocentric organisms, in which centromere activity is distributed along the entire chromosome, exhibit remarkable tolerance to chromosomal restructuring^2,11^. This architecture allows chromosomes fragmented by fission or joined by fusion while still retaining their ability to segregate properly^11^. The genus *Rhynchospora* (beak-sedges) is characterised by the specific association of its holocentromeres with *Tyba* satellite DNA arrays^5–7^ and large chromosome number variation^12,13^, providing a powerful model to investigate how karyotype evolution proceeds in the absence of centromere constraints.

Previous work from our lab has provided evidence that chromosomal fusions and, potentially, fissions underlie major karyotypic shifts in *Rhynchospora* and can be frequently associated with holocentromeric *Tyba* repeats^6^, suggesting that repetitive sequences may facilitate structural changes. In parallel, studies of meiotic recombination in *Rhynchospora breviuscula* revealed a striking departure from canonical monocentric patterns, as the crossover (CO) distribution does not associate with centromere distribution, gene density, epigenetic marks, or transposable elements^14^. Instead, the spatial patterning of meiotic recombination appears to be shaped by chromosome pairing dynamics, particularly telomere-led synapsis initiation, rather than sequence or chromatin state.

Karyotype alterations are expected to influence CO patterning^15,16^, yet how variation in chromosome number and structure shapes meiotic recombination remains unresolved. *Rhynchospora*, with its exceptional holocentric karyotype diversity (2*n* = 4 to 2*n* = 36) and tolerance to extensive fissions and fusions^12,13^, provides a natural framework in which chromosome-size shifts and structural rearrangements can be examined without the confounding lethality seen in monocentric systems. This genus, therefore, offers an unparalleled opportunity to test how recombination landscapes respond to karyotype evolution and to identify the structural principles linking chromosome architecture to CO output.

Here, we combined high-quality haplotype-phased chromosome-level genome assemblies, single-cell pollen sequencing, synapsis cytology, and chromatin conformation data across holocentric *Rhynchospora* species, encompassing 40 million years of evolution. By integrating structural and functional genomics, we show that chromosomal rearrangements frequently driven by *Tyba*-mediated fusions and fissions reconfigure recombination landscapes by altering chromosome size, chromatin loop architecture, and synapsis dynamics. Our results provide a mechanistic framework linking holocentric karyotype evolution to recombination plasticity, with broader implications for understanding how chromosomal structure shapes genetic diversity and speciation.

## Results

### The *Rhynchospora* pangenome reveals extensive holocentric karyotype evolution

To reconstruct the tempo and mode of karyotype evolution in holocentric *Rhynchospora*, we assembled chromosome-scale genomes for 17 *Rhynchospora* species, in addition to 3 previously available genomes^6,14,17^, yielding 56 haplotypes from 20 species spanning all major clades (**Supplementary Dataset 1 and Extended Data Table 1**). Gene-based synteny analysis^18^ recovered large conserved blocks across the pangenome, yet revealed pervasive structural reconfiguration among species. This comparative analysis reveals an unexpectedly rich diversity of karyotype-modifying processes, primarily driven by chromosome fissions and fusions, accompanied by frequent intrachromosomal inversions, whole-genome duplications (WGDs), allopolyploidy, and translocations, which occur at lower frequencies. Chromosome counts range from 2*n* = 4 (*R. tenuis*) to 2*n* = 36 (*R. rugosa*), underscoring the remarkable karyotype plasticity characteristic of holocentric chromosomes (**Fig. 1a, Extended Data Table 2**). The preservation of extensive collinearity despite widespread chromosome fissions and fusions indicates that speciation within *Rhynchospora* has proceeded primarily via rearrangements (i.e., chromosomal speciation) rather than rapid erosion of synteny.

**Fig. 1:**
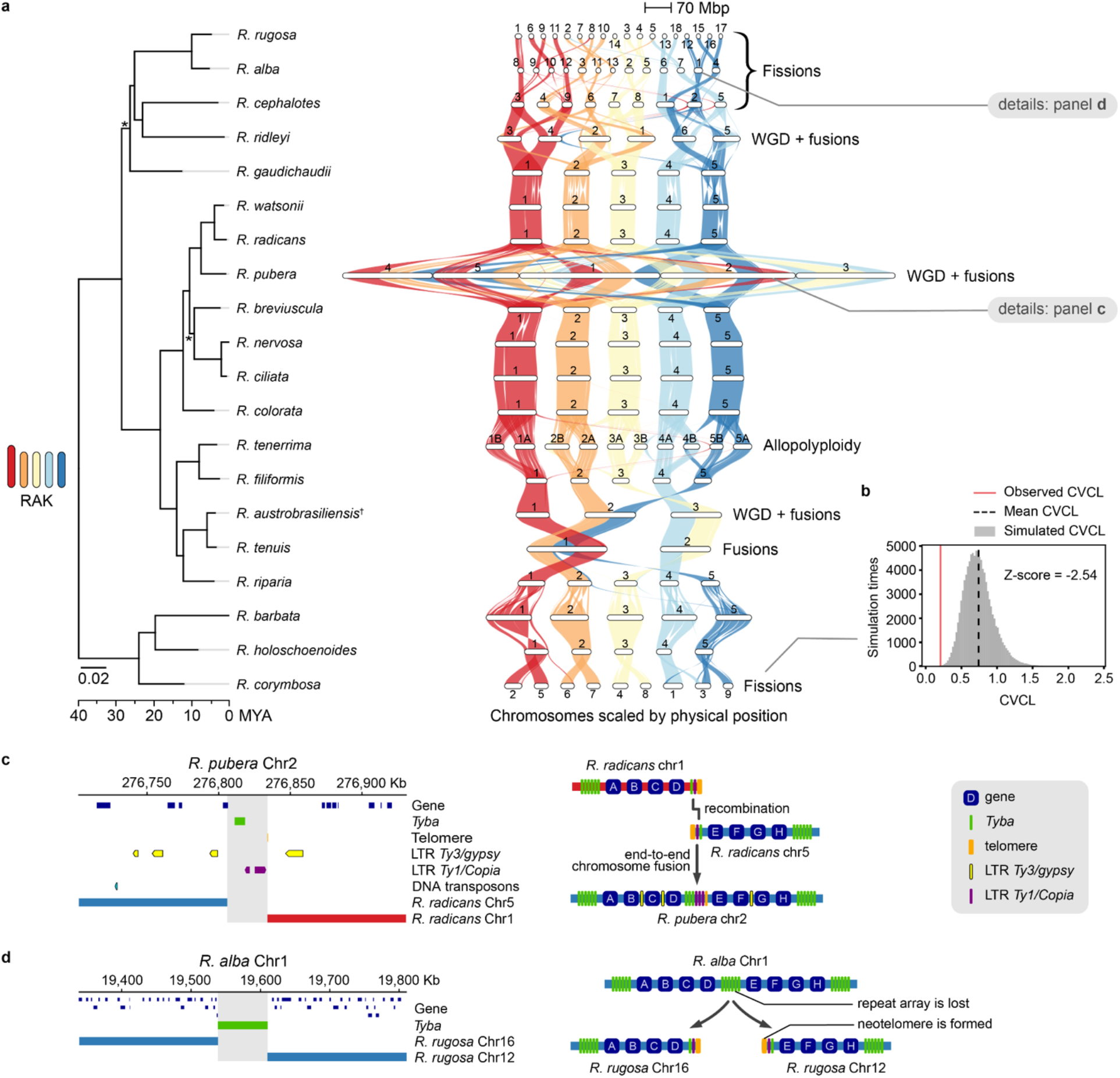
Karyotype evolution in holocentric *Rhynchospora* revealed by a comparative pangenome. (**a**) Time-calibrated orthoMCL phylogeny of 20 *Rhynchospora* species with chromosome-level haploid assemblies integrated with GENESPACE synteny. Conserved blocks are linked by coloured ribbons (haploid comparison shown) based on the most parsimonious *Rhynchospora* ancestral karyotype (RAK). Extensive interspecific re-wiring is explained primarily by fissions, fusions and inversions; while whole-genome duplications, allopolyploidy, and translocations occur at a lower frequency. Chromosome counts span 2*n* = 4 in *R. tenuis* to 2*n* = 36 in *R. rugosa*. Bootstrap values below 0.9 are indicated with asterisks. (**b**) Comparison between the observed coefficient of variation of chromosome length (CVCL) of *R. corymbosa* and the CVCL distribution of random simulations of chromosome breaking from *R. holoschoenoides* to *R. corymbosa*. A negative Z-score means that the dispersion of observed chromosome sizes is smaller than the dispersion of random chromosome breaking events, suggesting that the observed chromosome lengths of *R. corymbosa* are more balanced. (**c**) Representative fusion: *R. radicans* chromosomes 1 and 5 found in the origin of *R. pubera* chromosome 2; the fusion junction maps within *Tyba* repeats. (**d**) Representative fission: *R. alba* chromosome 1 corresponds to *R. rugosa* chromosomes 12 and 16; breakpoints coincide with centromeric *Tyba* arrays. Across the pangenome, 80 rearrangement junctions were identified, 58% overlapping *Tyba* arrays or LTR elements, implicating holocentromeric repeats as hotspots for both fusion and fission. **Extended Data Table 2** lists assemblies and counts; additional rearrangement examples are shown in **Supplementary Dataset 2**.

In *R. alba, R. rugosa*, and *R. cephalotes*, as well as independently in *R. corymbosa*, ancestral chromosomes split into smaller units (**Fig. 1a and Extended Data Fig. 1**), from the inferred ancestral *n* = 5 to *n* = 9–18 via fission rather than polyploidization. *R. rugosa* even formed the highest haploid chromosome number in the genus^13^. In contrast, chromosome fusion events are less abundant in the genus but are frequently associated with events of WGD (specifically in cases of autopolyploidy). Several WGDs were uncovered, notably often coupled with extensive chromosome fusions and secondary chromosome number reduction. In addition to the previously reported extreme case found in *R. pubera*, where two rounds of WGDs were followed by large-scale fusions that reduced its chromosome number to a complement with *n* = 5, a newly identified WGD in *R. ridleyi* shows an analogous trajectory, with fusion-mediated diploidization reducing the chromosome number to *n* = 6 (**Fig. 1a and Extended Data Fig. 2**). A distinct polyploidy event was detected in *R. gaudichaudii*, which carries a triploid genome exhibiting extensive haplotype differentiation and few reciprocal translocations (**Fig. 1a and Supplementary Fig. 1**). By contrast, *R. tenerrima* represents the first documented case of allopolyploidy in *Rhynchospora* in which the parental subgenomes remain largely collinear and free of major interchromosomal rearrangements (**Fig. 1a and Supplementary Fig. 2**). In *R. tenuis*, which carries the lowest chromosome number in the genus (*n = 2*), we previously found that its reduced chromosome number is derived from three chromosome fusions^6,17^. Now, we have revealed that its hypothesised intermediate structural state is preserved in the autohexaploid *R. austrobrasiliensis*, supporting a stepwise fusion pathway during chromosome number reduction in this lineage (**Fig. 1a and Supplementary Fig. 3**).

**Fig. 2:**
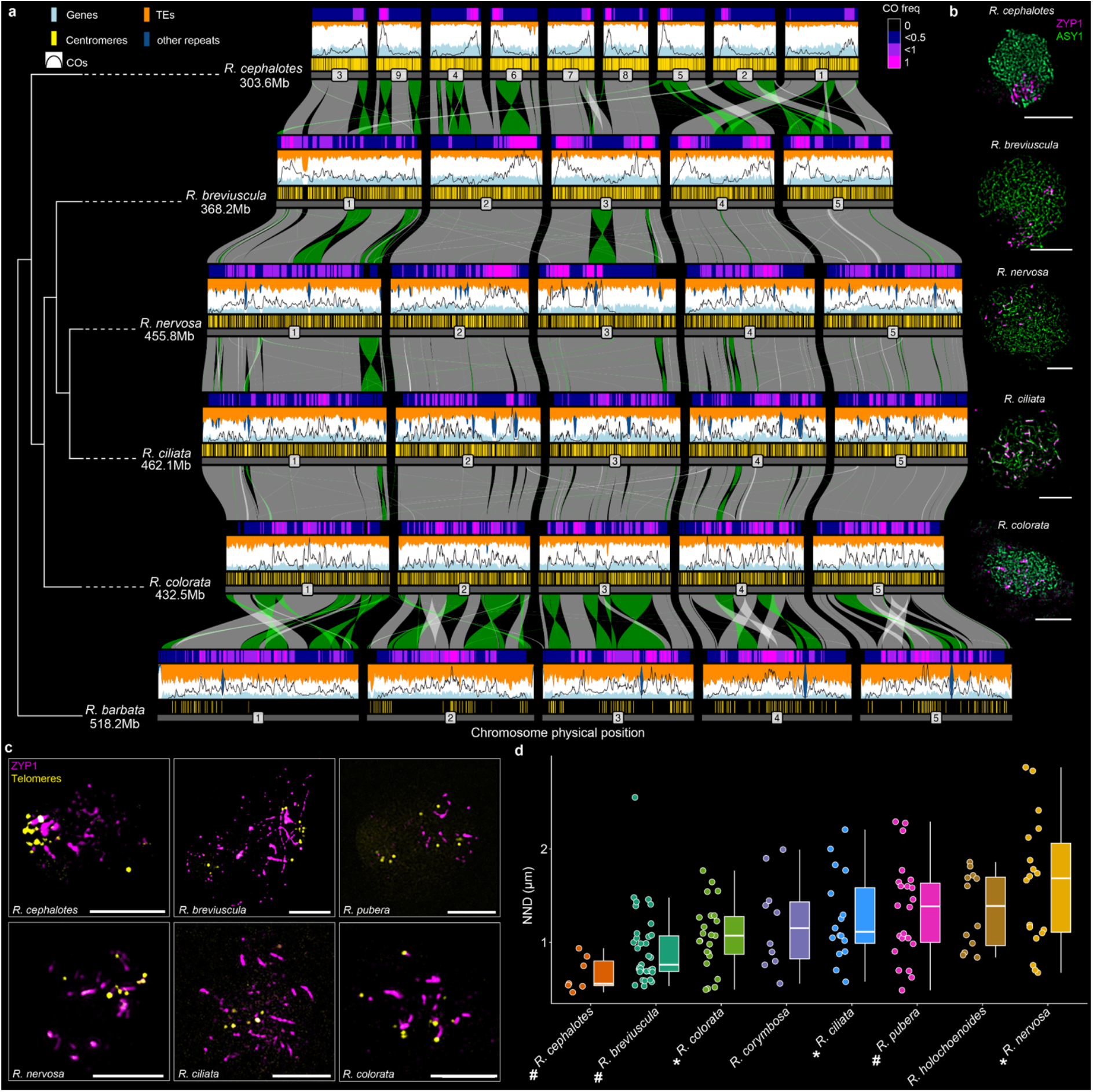
Synapsis organisation differs across *Rhynchospora* species and correlates with crossover landscape diversity. (**a**) Comparative synteny and CO maps derived from single-gamete sequencing across six *Rhynchospora* species, ordered by phylogeny (left). Chromosomes are shown scaled by physical length and aligned by conserved syntenic blocks (grey ribbons) and inversions (green ribbons). Tracks above each chromosome show gene density (light blue), transposable elements (orange), other repeats (dark blue), centromeric *Tyba* arrays (yellow), and CO positions (black lines). CO frequency is shown as a heatmap above the tracks. Species display either strongly distal-biased (i.e., *R. cephalotes* and *R. breviuscula*) or more irregular (i.e., *R. nervosa, R. ciliata, R. colorata* and *R. barbata*) CO distributions despite broadly conserved genomic features. To facilitate cross-species comparison, CO rates (cM/Mb) tracks were min–max (0-1) normalised per species; class assignment reflects consistent visual patterns. The minimum CO rates are 0 for all species, and the maximum CO rates are 9.25, 2.34, 2.28, 2.78, 2.22, 1.94 cM/Mb, following the order of the species on the phylogenetic tree from top to bottom. Across seven species, 29,433 COs were detected. **Extended Data Fig. 4** shows per-species detailed recombination maps; species lists and sample information are provided in **Extended Data Table 3**. (**b**) Immunofluorescence of ASY1 and ZYP1 during early zygotene reveals differential synapsis dynamics: species with distal-biased COs show clustered synapsis initiation, whereas species with irregular CO distributions show diffuse initiation of synapsis across chromosomes. Detailed images are shown in **Extended Data Fig. 5**. (**c**) High-resolution images of early meiotic prophase nuclei showing ZYP1 (magenta) and telomeres (yellow). In species with distal-biased recombination (for example, *R. cephalotes* and *R. breviuscula*), synapsis initiates in clustered regions near chromosome ends, whereas species with irregular CO landscapes (for example, *R. nervosa* and *R. colorata*) show a looser bouquet and dispersed synapsis initiation sites across the nucleus. Detailed images are shown in **Extended Data Fig. 6**. (**d**) Quantification of nearest-neighbour distances (NND) between telomere FISH signals across species. Boxplots show median, interquartile range, and distribution of NND values. Species with distal-biased CO landscapes (#) showed the smallest NNDs compared with species showing irregular recombination patterns (*). Normalised NND values are shown in **Supplementary Fig. 7**.

Across all species, chromosome sizes remain remarkably similar within each karyotype despite extensive histories of chromosome fissions and fusions (**Fig. 1a**). To quantitatively assess the dispersion of chromosome sizes across *Rhynchospora*, we calculated the coefficient of variation of chromosome length (CVCL) for species that experienced substantial karyotype restructuring. In all such cases, observed CVCL values were significantly lower than the expectation of chromosome-size distributions under random chromosome breaks or fusions (**Fig. 1b and Extended Data Fig. 3**). These results demonstrate that structural rearrangements consistently generate chromosome complements with near-even size distributions, a recurrent feature of holocentric karyotypes^13,19–21^. Together, our findings indicate strong evolutionary constraints favouring interchromosomal size symmetry in holocentric genomes.

**Fig. 3:**
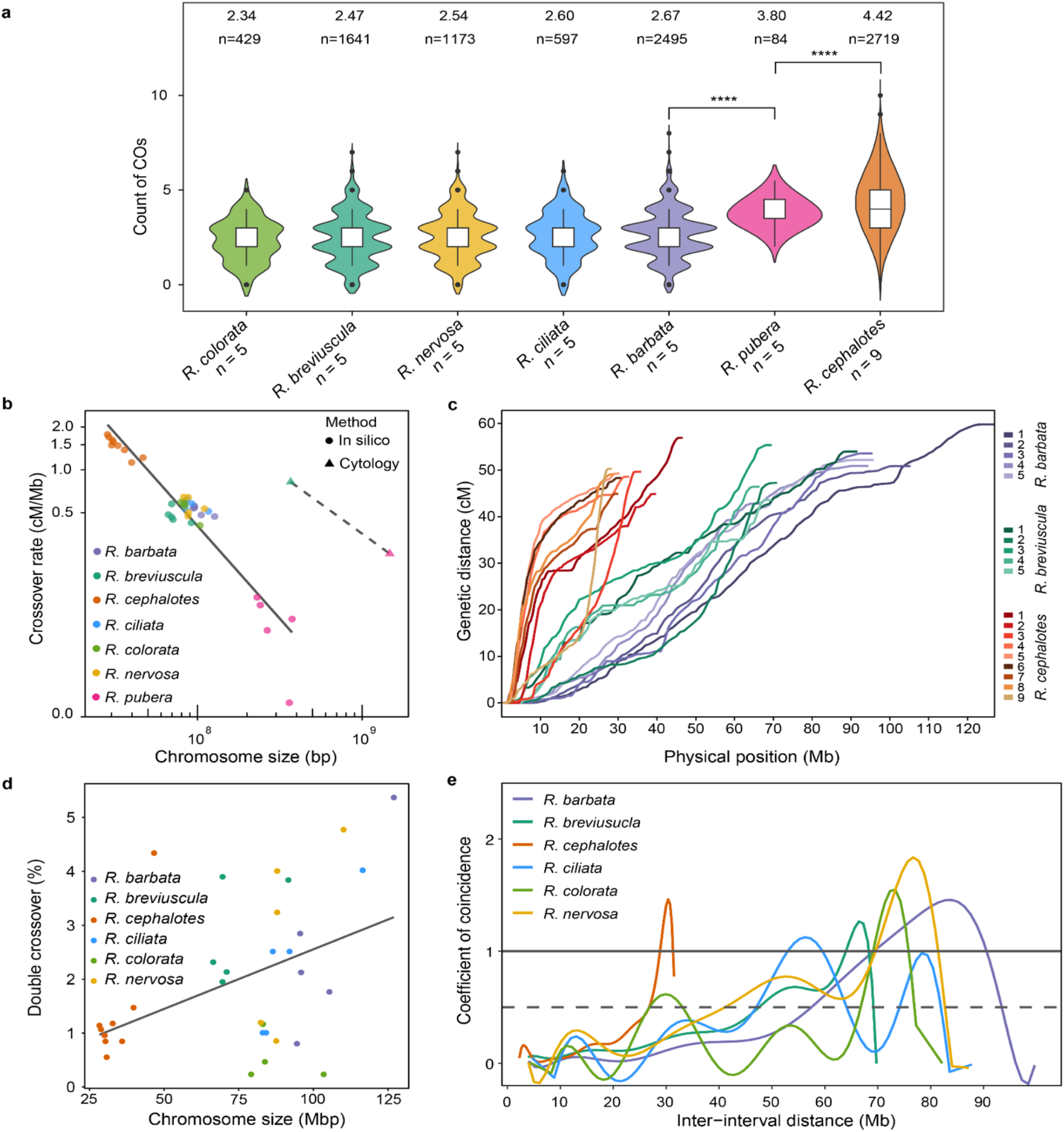
Chromosome number and size quantitatively govern crossover output in holocentric *Rhynchospora*. (**a**) Total CO counts across seven *Rhynchospora* species ordered by increasing mean CO count. Species with higher chromosome numbers show increased CO counts, consistent with CO assurance. CO numbers for *R. pubera* were estimated from MLH1 cytological counts and normalised for comparison with pollen sequencing–based estimates in other species. Haploid chromosome numbers (*n*) are given below each species name. Shapiro test for all species indicates non-normal distribution. Two-sided Mann-Whitney U test between *R. barbata* and *R. pubera*: *p* < 2.2e-16; between *R. pubera* and *R. cephalotes*: *p*=3.13e-05. Other neighboring pairs have no significant difference. (**b**) Inverse relationship between chromosome physical length and recombination rate (cM/Mb) across all chromosomes and species. Smaller chromosomes exhibit higher CO density than larger chromosomes (*In silico* method: adjusted *R*^2^ = 0.89, *p* < 2.2 × 10^−16^). (**c**) Marey maps for representative species illustrating how recombination rate scales with chromosome size and landscape class. Small chromosomes show steep genetic–physical distance relationships, whereas larger chromosomes display flatter or segmental profiles depending on CO distribution mode. Further per species detailed Marey maps are shown in **Supplementary Fig. 8**. (**d**) Frequency of double COs plotted against physical chromosome length, showing increased double CO occurrence on longer chromosomes (adjusted *R*^2^ = 0.17, *p* =0.0096). (**e**) Coefficient of coincidence as a function of inter-interval distance across species, indicating chromosome-size-dependent modulation of crossover interference. CoC<1 indicates CO interference within this interval distance. Together, these analyses demonstrate that chromosome number and physical size are major determinants of crossover number, density, and higher-order organisation in holocentric genomes.

Remarkably, across these diverse karyotypes, all major clades retain species with the same five ancestral syntenic blocks, which are preserved with remarkable stability despite extensive karyotype remodelling. These blocks undergo frequent intrachromosomal inversions but only rare interchromosomal translocations. The pervasive recurrence of *n* = 5 across phylogenetically distant lineages, and its clear correspondence to five conserved chromosomal synteny units in **Fig. 1a**, supports *n* = 5 as the most likely *Rhynchospora* ancestral karyotype (RAK).

Our comparative analysis enables precise reconstruction of rearrangement histories. We have shown that the *Rhynchospora*-specific holocentromeric *Tyba* repeat is frequently associated with chromosome fusion events^6^. For example, a fusion between chromosomes 1 and 5 in *R. radicans* is found in chromosome 2 of *R. pubera* with a functional *Tyba* array being found precisely at the rearrangement point (**Fig. 1c**). Remarkably, we have now found that also fission events, for example, a fission of *R. alba* chromosome 1 accounting for the origin of *R. rugosa* chromosomes 12 and 16 is precisely mapped to a *Tyba* array in the *R. alba* chromosome (**Fig. 1d**). In both cases, the inferred breakpoints localise within centromeric *Tyba* arrays, implicating repeat-rich holocentromeric regions in mediating both breakage and joining.

To systematically test this association, we applied a breakpoint-detection pipeline (see **Methods**) and mapped 80 independent junctions (rearrangement regions) across the pangenome, from which 69 were classified as fusion or fission events (**Extended Data Table 2, Supplementary Dataset 2**). Comparative analyses with closely related species and outgroups to infer directionality showed that 58% of breakpoints overlap *Tyba* arrays or transposable elements (**Fig. 1c–d, Supplementary Dataset 2, Extended Data Table 2**). Our breakpoint analysis shows that *Tyba* arrays frequently overlap fusion and fission junctions, consistent with a model for ectopic recombination and/or defective double-strand-break repair in *Tyba*-rich chromatin^6^. These findings further provide a mechanistic basis for the chromosome-number shifts observed across *Rhynchospora*.

### Divergent recombination landscapes correlate with synapsis dynamics

To assess how the dynamic karyotype variation found in *Rhynchospora* impacts meiotic recombination, we generated genome-wide CO maps for five additional *Rhynchospora* species (*R. barbata, R. cephalotes, R. ciliata, R. colorata*, and *R. nervosa*) using single-cell sequencing of pollen nuclei and integrated these with a genetic map from a *R. pubera* F_2_ population (**Extended Data Table 3; Supplementary Fig. 4**) together with the available *R. breviuscula* CO map^14^.

We detected 29,433 COs across the seven species and found that recombination landscapes broadly group into two apparent modes (**Fig. 2a, Extended Data Fig. 4, Supplementary Fig. 5**): a strongly distal-biased pattern, with COs enriched at one or both chromosome ends (*R. cephalotes, R. breviuscula, R. pubera*), and an irregular pattern with COs distributed more evenly along each chromosome (*R. barbata, R. ciliata, R. colorata, R. nervosa*). Notably, distal enrichment was observed irrespective of chromosome number and size, e.g., *R. cephalotes* (*n* = 9, mean chromosome size 33.7 Mbp) and *R. breviuscula* (*n* = 5, mean chromosome size 74 Mbp). Together, these patterns indicate that CO spatial distributions are shaped by mechanisms that operate independently of chromosome size or number.

**Fig. 4:**
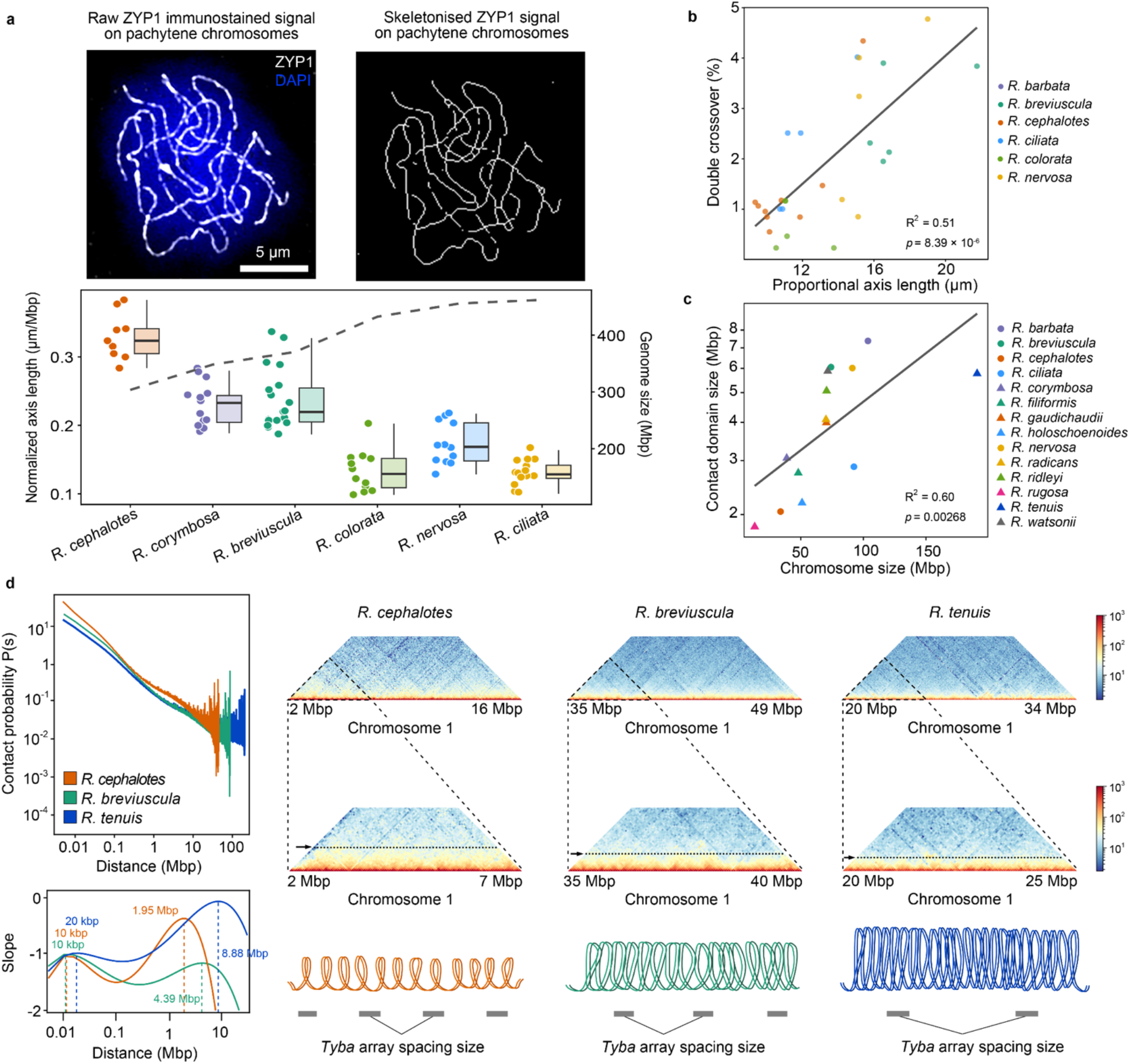
Chromatin loop geometry links chromosome size to axis length and crossover frequency in holocentric *Rhynchospora*. (**a**) Raw ZYP1 immunofluorescence on *R. breviuscula* pachytene chromosomes used to measure synaptonemal complex axes (left). Skeletonised ZYP1 signal illustrating axis tracing and quantification per chromosome (right). Quantification of axis length per megabase was measured in pachytene stage across species (bottom). The species was ordered by their genome sizes, indicated by the dashed line. Detailed and additional representative images are shown in **Extended Data Fig. 7**. (**b**) Frequency of double COs plotted against proportional chromosome axis length, showing increased double CO occurrence on longer chromosome axis. Linear regression shows a positive correlation between the frequency of double COs (DCOs) and axis length (adjusted *R*^2^ = 0.51, *p* = 8.39 × 10^−6^). (**c**) Comparative analysis of the loop size measurement and the average chromosome sizes across 14 *Rhynchospora* species that have deep sequenced Hi-C. Linear regression shows a positive correlation between the mean chromosome size and log10 scale of the chromatin contact domain size (adjusted *R*^2^=0.60, *p*=0.00268). The contact domain size is defined by the first turning point of contact probability P(s) slope curve (**Supplementary Dataset 4**). (**d**) Comparative analysis of chromatin conformation from somatic Hi-C data from three representative *Rhynchospora* species with different chromosome sizes to illustrate the effect of chromosome size on loop size architecture, hinting at a possible role of the spacing dynamics of between consecutive holocentromeric *Tyba* repeat arrays in overall chromosomal axis length^7^.

To further understand the link between sequence collinearity and meiotic recombination frequencies, we developed a plotting strategy to integrate DEEPSPACE synteny maps with CO maps (**Fig. 2a**). Despite extensive conservation of synteny across species, syntenic intervals frequently display divergent recombination profiles (**Fig. 2a**), indicating that CO patterning is not conserved with sequence collinearity. For example, the centre of chromosome 2 in *R. breviuscula* shows low recombination. In contrast, its syntenic counterpart resides at a chromosome end in *R. cephalotes* and exhibits a pronounced recombination peak (**Fig. 2a**), indicating that chromosome-scale dynamics, rather than sequence collinearity, shape recombination.

On a broader scale, CO location showed no correlation with canonical genomic features (**Fig. 2a and Extended Data Fig. 4, Supplementary Dataset 3**). CO density did not track gene content, transposable elements, *Tyba* repeat arrays, or single-nucleotide polymorphisms (SNPs). This contrasts with monocentric species, where centromere-proximal and heterochromatic regions typically suppress recombination^15^, and indicates that CO patterning in *Rhynchospora* is largely decoupled from local sequence or epigenetic state likely due to its decompartmentalised genome organisation.

To investigate the mechanistic basis underlying the divergent recombination landscapes observed, we examined meiotic chromosome behaviour by high-resolution immunofluorescence of the axial element ASY1 and the SC transverse filament ZYP1. Species with distal-biased CO landscapes (*R. breviuscula, R. cephalotes, R. pubera*) displayed clustered synapsis initiation. In contrast, species with COs irregularly distributed (*R. nervosa, R. ciliata, R. colorata*) exhibited more loose ZYP1 signals at the onset of synapsis initiation (**Fig. 2b and Extended Data Fig. 5**). We also investigated synapsis initiation in species lacking recombination maps due to low heterozygosity. *R. holoschoenoides* showed clustered synapsis, whereas *R. corymbosa* showed non-preferential early synapsis (**Extended Data Fig. 5**), suggesting corresponding divergence in recombination patterning.

**Fig. 5:**
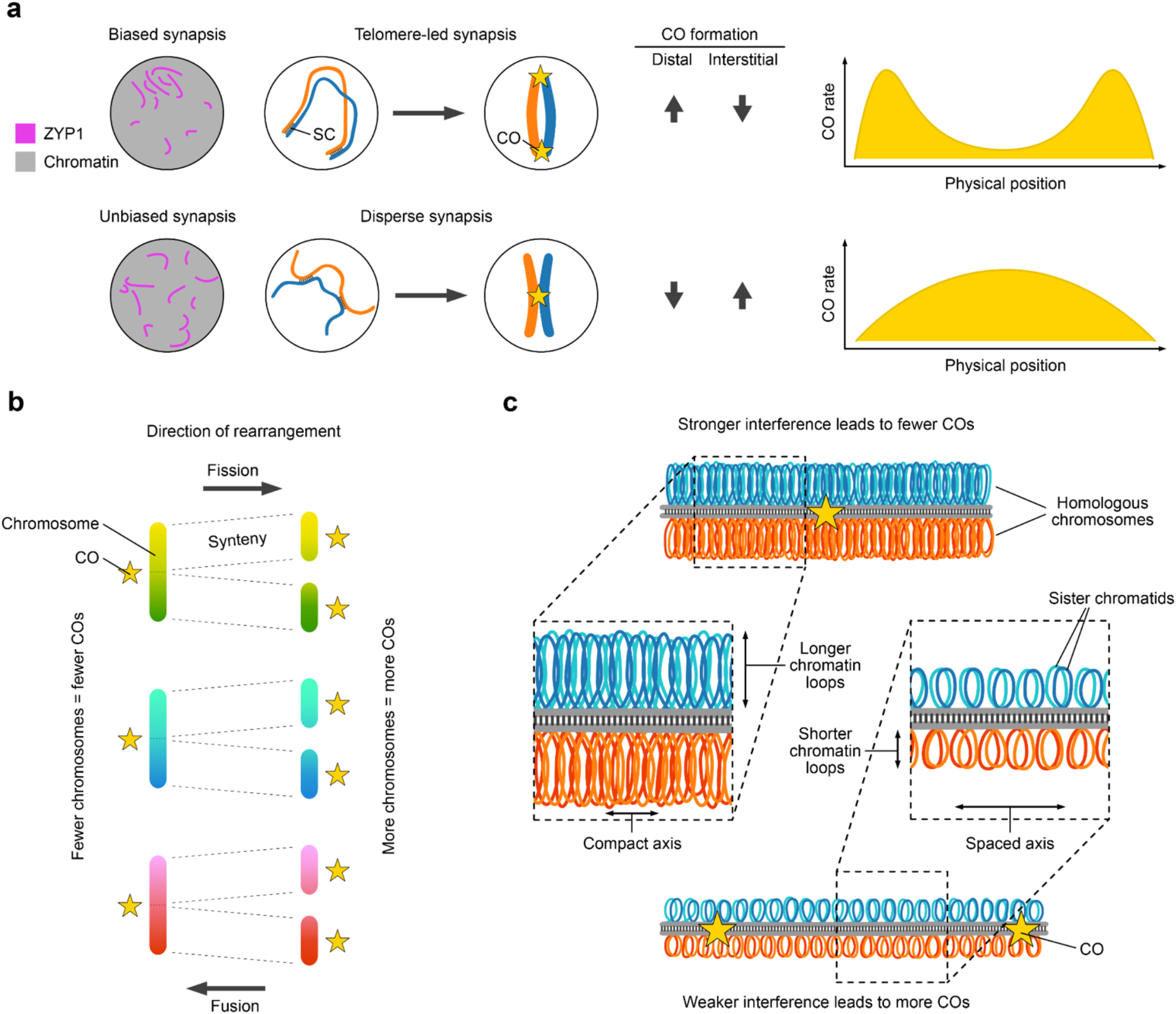
A structural-mechanistic framework linking holocentric karyotype evolution to crossover patterning in *Rhynchospora*. (**a**) Synapsis modes determine CO spatial patterning: clustered, telomere-led synapsis (biased) generates distal CO enrichment and terminal ramps in CO rate along physical position; dispersed synapsis (unbiased) yields more interstitial or irregular CO placement. (**b**) Effects of chromosome number on total COs: fission increases chromosome number and elevates total COs via CO assurance; fusion lowers chromosome number and reduces total COs (schematic chromosomes and SC, numbers illustrate expected differences in CO counts). (**c**) Loop-axis geometry sets CO capacity and interference: large chromosomes with longer loops and a compact axis show stronger interference and fewer COs; small chromosomes with shorter loops and an expanded axis show weaker interference and more COs. Together, these panels synthesise how chromosome fissions and fusions, synapsis dynamics, and loop-axis organisation jointly control both the quantitative output and spatial distribution of COs in holocentric *Rhynchospora*.

To link synapsis initiation with possible differences in telomere bouquet, we performed immuno-FISH with ZYP1 and telomere FISH on early zygotene cells. We then analysed the bidimensional distribution of telomeres to measure their clustering during zygotene. Interestingly, *R. cephalotes, R. breviuscula* and *R. pubera* show a tighter clustering of telomeres consistent with a pronounced telomere bouquet, followed by synaptonemal complex elongation from chromosome ends. In contrast, *R. nervosa, R. ciliata* and *R. colorata* appear to have a less pronounced telomere bouquet configuration consistent with synapsis initiating from multiple spots (**Fig. 2c–d, Extended Data Fig. 6, Supplementary Fig. 6a**). Together, these data support a correlation between synapsis dynamics and CO spatial distributions, implicating early telomere-led pairing and subsequent synapsis progression as primary determinants of recombination landscapes in holocentric *Rhynchospora*, largely independent of underlying genomic features.

### Chromosome size governs crossover frequency

We next asked how karyotype configuration influences the number and density of meiotic COs. Because CO assurance, which mandates at least one CO per chromosome to ensure proper segregation^22^, it is expected that a higher chromosome number also exhibits higher total CO numbers. Indeed, *R. cephalotes* (*n* = 9), with an elevated chromosome count, exhibits a higher total CO count than species with fewer chromosomes, e.g., *R. breviuscula* (*n* = 5). Across species, chromosome number was positively correlated with total CO number (**Fig. 3a**), indicating that changes in chromosome count directly scale recombination output.

We then quantified the relationship between physical length and recombination rate. Pooling all chromosomes from the seven *Rhynchospora* species with available CO maps, we observed a strong inverse relationship between chromosome length and CO density, measured as COs per megabase (linear model of log10(CO rate) ~ log10(chromosome length): adjusted R^2^ = 0.89, *p* < 2.2 × 10^-16^, **Fig. 3b**). Thus, smaller chromosomes carry proportionally more COs per unit of DNA than larger chromosomes. This size dependence is consistent with structural constraints: assurance imposes at least one CO even on short chromosomes, elevating per-megabase rates, and shorter physical lengths likely increase the effective axial substrate per megabase available for CO designation. The relationship holds across karyotypes with different rearrangement scenarios; for instance, *R. pubera* is characterised by several chromosome fusions, while *R. cephalotes* is derived mostly by fissions.

Marey maps further illustrate how recombination scales with chromosome size (**Fig. 3c and Supplementary Fig. 8**). In *R. cephalotes*, a species with a distal-biased CO landscape, small chromosomes exhibit steep, segmental cM–Mb relationships, with pronounced slopes at one or both ends. Species with irregular landscapes (e.g., *R. barbata*) display near-linear slopes along chromosome arms, consistent with diffused CO placement. Together, these trends indicate that karyotype evolution quantitatively modulates recombination through changes in chromosome number and size.

Finally, we asked whether chromosome size also influences higher-order recombination features, including the frequency of double COs (DCOs) and the strength of CO interference^23^, where the occurrence of one CO suppresses the formation of another nearby. Thus, interference reduces the frequency of DCOs relative to random expectations. We found that short chromosomes exhibit a lower proportion of DCOs, whereas longer chromosomes show increased DCO frequency (linear model: adjusted *R*^2^ = 0.17, *p* = 0.0096, **Fig. 3d**), consistent with their greater physical length and expanded opportunity to accommodate multiple COs. CO Interference analyses using the coefficient of coincidence formula (**see Methods**) further reveal that interference strength scales with chromosome size across species (**Fig. 3e**), linking physical chromosome architecture to the regulation of CO spacing. Together, these results show chromosome size as a major determinant of CO frequency^24^, while also shaping the higher-order organisation of recombination through its effects on CO interference.

### Chromosome and chromatin loop size link axis length to crossover frequency

To uncover the mechanism underlying the inverse relationship between chromosome size and recombination rate, we quantified meiotic axis architecture and chromatin loop geometry across *Rhynchospora*. We first measured synaptonemal complex (SC) axes by immunostaining ZYP1 on pachytene chromosomes and skeletonising the signal to extract axis length per bivalent and per chromosome (**Fig. 4a and Supplementary Figure 6b**). Across species and chromosome classes, smaller genomes exhibited greater axis length per megabase, whereas larger genomes showed reduced axis length per DNA unit (**Fig. 4a and Extended Data Fig. 7**).

Next, we asked whether meiotic chromosome axis length influences the frequency of DCOs. Variation in DCO frequency may reflect underlying chromosome properties. Ongoing research and theoretical models propose that CO interference arises from processes operating along the chromosomal axis, including HEI10 diffusion and aggregation along the synaptonemal complex (coarsening model)^25,26^, as well as chromosome-axis-based mechanical stress propagation (beam-film model)^27^. Both models proposed a central role of meiotic chromosome axes. Consistent with these predictions, we measured meiotic chromosome axis lengths in *Rhynchospora* species having CO counting and found that chromosomes with longer axes exhibit a higher probability of DCOs (linear model: adjusted *R*^2^ = 0.51, *p* = 8.39 × 10^−6^, **Fig. 4b**). Notably, this association was substantially stronger than that observed when DCO probability was correlated with physical chromosome length measured in base pairs (**Fig. 3d**). These observations suggest that meiotic chromosome compaction and axis organization, rather than genomic size alone, are key to CO frequency.

Because axis length is inversely related to chromatin loop size^28–31^, we next estimated average chromatin loop size from Hi-C contact decay profiles as a proxy for loop geometry. Chromosome size showed a clear positive correlation with chromatin contact domain size (Adjusted *R*^2^=0.60, *p*=0.00268): larger chromosomes tended to be folded into larger domain sizes, while smaller chromosomes were organised into smaller ones (**Fig. 4c**). Although our Hi-C data was generated from non-meiotic tissue, we reason that relative differences in chromatin architecture across chromosomes are likely conserved across different tissues and cell-types within an individual, including meiosis, allowing loop size to serve as a meaningful proxy for axial organisation^32,33^.

The correlation between chromosome size and chromatin domain sizes estimated from Hi-C contact-decay profiles is consistent with the proposed mechanistic role of the holocentromeric *Tyba* repeat on the architecture of *Rhynchospora* holocentric chromosomes supported by polymer modeling^7^. Consistent with this view, we modeled the effect of *Tyba* array spacing on the overall axis length of *Rhynchospora* chromosomes and compared with real measurements. Indeed, we observed a close match between modeled axis length and our measurements (*r*(6)=0.946), confirming that loop length is inversely related to axis length (*r*(6)=-0.963 for simulated axis length and *r*(6)=-0.851 for experimentally measured axis length; **Supplementary Fig. 9**).

These findings match the prediction that increasing *Tyba* inter-array spacing expands chromatin loops while tighter spacing constrains them^7,34,35^ (**Fig. 4d**). Through this mechanism, small chromosomes with short loops and extended axis length per megabase present more potential sites for CO designation, resulting in elevated CO density. In contrast, large chromosomes, with long loops and relatively compressed axes, limit access for recombination machinery and suppress CO formation. This loop-axis mechanism operates alongside CO assurance to scale recombination with chromosome number, providing a coherent framework in which karyotype evolution links holocentromere architecture, chromosome size, and the quantitative reshaping of recombination landscapes in holocentric *Rhynchospora*.

## Discussion

Our comparative analysis across the genus *Rhynchospora* shows that holocentric chromosome architecture permits exceptional tolerance to both fissions and fusions. Large syntenic blocks are retained despite extensive karyotype restructuring, indicating that speciation proceeds predominantly through rearrangements rather than erosion of sequence collinearity^9,36^. We identify the holocentromeric *Tyba* satellite repeat as the primary hotspot for both fusion and fission events, consistent with repeat-rich regions acting as fragile sites^6,37^ that catalyse structural change while being protected from lethality through distributed kinetochore activity. These properties help explain the striking chromosome-number volatility in *Rhynchospora* and parallel patterns described in other holocentric groups^38,39^.

Holocentric species also tend to maintain highly symmetric karyotypes, with chromosomes of similar physical length. This pattern is widespread across plants, nematodes and insects, suggesting that chromosome-size uniformity is not incidental but reflects underlying evolutionary pressure^36,38– 41^. In *Rhynchospora*, near-equal chromosome sizes persist despite extensive histories of fission and fusion. A comparable pattern occurs in the Atlas blue butterfly (*Polyommatus atlantica*), where 227 autosomes produced by fragmentation of 24 ancestral chromosomes remain uniformly sized^19^. A mechanistic explanation is that holocentric chromosomes assemble kinetochore activity along their entire length, such that chromosome size sets total kinetochore mass^42,43^. A chromosome fission that generates a fragment with an uneven kinetochore pool relative to the other chromosomes would disrupt spindle force balance, increasing the risk of mis-segregation^44–47^. Further chromosome fissions or fusions would help restore kinetochore-size parity. However, chromosome fissions may occur more frequently than fusions, as they require a single DSB followed by end-healing. In contrast, fusions require two DSBs plus an ectopic repair event^4^, making consecutive fissions more likely during karyotype evolution in holocentric groups.

Furthermore, changes in chromosome length could facilitate meiotic/holokinetic drive^48^, which, in turn, could fix a new karyotype after only one (or a few) plant generations. Meiotic drive mechanisms in holocentric organisms have been documented in wood white butterfly hybrids (*Leptidea sinapis*), where female drive may favour shorter chromosomes arising by fissions^49^, and more recently in *Rhynchopsora tenuis*, where we show that simultaneous male and female meiotic drive counteract each other, keeping a highly asymmetric karyotype stable across generations^17^.

The structural plasticity of *Rhynchospora* chromosomes also has clear consequences for meiotic recombination. Across the genus, CO landscapes fall into two distinct classes, one strongly distal-biased and the other irregular, and these patterns show little association with local genomic features^14^. Instead, cytology links CO placement to synapsis dynamics, in which species with more distal-biased COs exhibit clustered synapsis initiation and irregular species exhibit dispersed initiation (**Fig. 5a**). The mechanism underlying this phenomenon remains elusive. Previously, we proposed that distally positioned recombination intermediates might have a temporal advantage at being processed into final COs where synapsis is imposed early^14^. These conclusions are consistent with findings in monocentric mammals, where chromosomal fusions have been shown to remodel 3D genome folding in the germ line and, as a consequence, alter recombination landscapes independently of local sequence composition^10^, highlighting chromosome architecture as a conserved regulator of meiotic recombination across eukaryotes.

Across species, total CO number scales positively with chromosome number, consistent with CO assurance^50^ (**Fig. 5b**), whereas CO density scales inversely with chromosome length. The chromatin loop-axis framework readily explains these observations. During meiosis, axis length and loop size are inversely correlated^28–30,51–53^. Short chromosomes, therefore, have a longer axis per unit DNA and more COs, while long chromosomes have a shorter axis and fewer COs^30,54^. CO interference^55,56^ likely reinforces these differences. While absolute loop sizes likely differ between mitotic and meiotic contexts^57^, relative differences are expected to persist^33^, supporting loop length as a meaningful proxy for recombination potential. Together with species-specific synapsis modes, the chromatin loop-axis organisation can explain the quantitative scaling and spatial diversification of recombination across the genus (**Fig. 5c**).

An important insight from our analysis is that chromosome size contributes substantially to variation in recombination frequency^24,40^, acting through a structural hierarchy in which chromosome length influences chromatin-loop size^28,33^, and, in turn, the extent of meiotic axis available for CO formation, with this relationship being particularly evident for the frequency of double COs. Because shorter chromosomes are organised into smaller loops and therefore possess a proportionally longer axis per megabase, they provide a larger substrate for DSB formation and CO designation. In contrast, longer chromosomes, with larger loops and a more compact axis, inherently suppress recombination^30,51^. At the same time, a recent work in mice indicates that chromosome and axis length are not sufficient on their own to predict CO output, as species- and context-specific regulation of crossover designation efficiency, interference strength, and pathway usage can modulate recombination outcomes within the structural constraints imposed by chromosome architecture^58^.

Together *Rhynchospora* offers a uniquely powerful system for resolving this relationship, as its holocentric chromosomes tolerate extensive fissions and fusions that generate broad, naturally occurring variation in chromosome size while preserving synteny. Moreover, its repeat-based holocentromeres, composed of *Tyba* arrays, influence chromatin-loop spacing and thus directly contribute to loop-axis geometry^7^. Together, these features position *Rhynchospora* as an ideal model for dissecting how chromosome architecture influences meiotic recombination, and for establishing general principles through which karyotype evolution shapes the recombination landscape across eukaryotes.

## Methods

### Plant Material

Plants from naturally occurring populations of the species listed in **Extended Data Table 1**, growing in Brazil, were collected and subsequently cultivated under controlled greenhouse conditions (16 h daylight, 26 °C, >70% humidity).

### DNA isolation

High-molecular-weight DNA from the species listed in **Extended Data Table 1** was isolated from 1.5 g of material using the NucleoBond HMW DNA kit (Macherey-Nagel). Quality was assessed with a FEMTOpulse device (Agilent), and quantity was measured with a Qubit fluorometer (Thermo).

### PacBio

HiFi libraries were prepared according to the manual for “SMRTbell Express Template Prep Kit 2.0” or the “SMRTbell® prep kit 3.0”, with an initial DNA fragmentation by Megaruptor-3 (Covaris) and final library size selection by BluePippin (Sage Science). The size distribution was again controlled using FEMTOpulse (Agilent). Size-selected libraries were then sequenced on a Sequel IIe device with Binding kit 2.0 and Sequel II Sequencing Kit 2.0 for 30 h (*Pacific Biosciences*).

### Arima Hi-C

Plant tissues were cross-linked with 1% formaldehyde for 30 minutes at room temperature, and the reaction was quenched with 125 mM glycine for 10 minutes. Subsequently, the tissues were ground in a TissueLyser at 30 Hz for 3 minutes. Nuclei extraction was performed using the CelLytic PN Plant Nuclei Isolation/Extraction Kit (Sigma-Aldrich, Burlington, MA, USA) according to the manufacturer’s protocol. Hi-C libraries were prepared using the Arima High Coverage Hi-C Kit (Arima Genomics, A410110, Carlsbad, CA, USA) following the manufacturer’s instructions, and were then sequenced paired-end (2 × 150 bp) on a NextSeq 2000 instrument (Illumina, San Diego, CA, USA).

### Omni-C

For each species, a single chromatin-capture library was prepared from 0.5 g of fresh-weight material input. All treatments were administered according to the kit vendor’s recommendations for plants (Omni-C, Dovetail). As a final step, an Illumina-compatible library was prepared (Dovetail) and paired-end, 2 x 150 bp, deep-sequenced on a NextSeq 2000 (Illumina).

### Genome assemblies and scaffolding

Genome sequencing and assembly: PacBio HiFi reads were generated for 20 *Rhynchospora* species and assembled with Hifiasm with default settings. Hi-C sequences were included when running HiFiasm^59^ when available. Scaffolding was performed by aligning Hi-C reads to contigs from HiFiasm using Juicer and 3D-DNA^60,61^ for homozygous genomes. Alternatively, phased scaffolding was performed using HapHiC^62^ for heterozygous species. Scaffolding of the triploid genome of *R. gaudichaudii* was done using CPhasing^63^.

### K-mer-based genome size and heterozygosity estimation

Genome size and heterozygosity levels were estimated by GenomeScope2.0^64^ *K*-mer counting was performed using jellyfish v2.3.1^65^ with PacBio HiFi reads, which were used for genome assembly, as the input. Kmer size was set to 21. The resulting binary output (.hist) from jellyfish was then converted to a text histogram file. The histogram was then analysed with GenomeScope2.0 using k=21 and ploidy=2, except for *R. austrobrasiliensis*, where ploidy = 6 and *R. tenerrima*, where ploidy = 4.

### Repeat annotation

Repeat annotation was performed with DANTE and DANTE-LTR^66^ and Earl Grey v7.0.1^67^ using the respective plant TE library. *Rhynchospora*-specific holocentromeric *Tyba* repeat annotations were obtained from Wlodzimierz et al.^7^.

### Synteny analysis

Synteny analysis shown in **Fig. 1a** was performed using GENESPACE^18^ for 20 sets of single haplotypes from 20 *Rhynchospora* species.

Synteny analysis for comparative recombination maps in **Fig. 2a** was performed with DEEPSPACE (https://github.com/jtlovell/DEEPSPACE). Input data (assemblies, gene and repeat annotations and crossovers) were parsed and read into R with standard input methods. Syntenic blocks were inferred from windowed alignments with DEEPSPACE. In short, the query genome (largest of the pair) was split into 200020 non-overlapping evenly distributed 500bp windows and aligned to the target (smallest of the pair) with minimap2 with the following parameters: -r500,5000 -O6,26 -e2,1 -B4 -f0.001 -p0.75 -N4 -k23 -w15 -g1000 -F5000 -t8 -A1 -U50,500 --no-long-join --rmq=no -n1 -m0 --frag=yes. The windowed paf was parsed to syntenic hits with MCScanX_h (https://github.com/wyp1125/MCScanX), with gap and size parameters set to 5 hits. The resulting collinear hits were used as anchors to pull any hits within 5 rank-order positions, repruned to collinear hits and clustered with dbscan using eps radius of 2.6 positions and a minimum cluster size of 5. Adjacent hits more than 1Mb apart were split into separate blocks. Sliding windows were applied to gene and repeat density in 100kb overlapping 1Mb windows.

Crossovers were analyzed in two ways. First, a heatmap was built binning regions into four groups: none (black) = 0 COs, low (darkblue) = <0.5 COs, mid (purple) = <1 COs, high (magenta) >= 1COs. The CO rates of each species were normalized to a 0-1 scale based on their minimum and maximum CO rates. The minimum CO rates are 0 for all species, and the maximum CO rates are 9.25, 2.34, 2.28, 2.78, 2.22, 1.94 cM/Mb. All area plots have the same scale (0-1).

### Identification of break intervals in chromosome fusion and fission events

Whole genome alignments between different *Rhynchospora* species were conducted by minimap2 with ‘-cx asm10’. In both chromosome fusion and fission events, the larger chromosomes were taken as the target, and the smaller chromosomes were the query. Query chromosomes that account for no less than 1Mbp of the target chromosome length were the main components of the target. Each query chromosome ID was assigned to an integer. A sliding window with a 50kb window size and a 10kb step size was applied to each target chromosome to derive the query chromosome in this window by calculating the mean of the integer ID of the aligned query. The smoothed query integer ID was then input to a median filter for further denoising. The rough break intervals were defined at windows where consecutive non-zero first derivatives were found. To refine the break intervals by the last aligned locus of the query sequence, a binary signal partition approach was adopted. The complete pipeline is available on our project GitHub page: https://github.com/Raina-M/HoloRecomb_project. A chromosome is considered to have N fission events when it is composed by N+1 different query chromosomes. Only when the query chromosomes contain the broken target sequences and the joining of sequences happened at the end of the query chromosomes, a fusion event is counted (**Extended Data Table 2**). If a query chromosome has the same broken target fragments multiple times, only one fusion event is considered related to this target chromosome.

### Random simulation of chromosome fissions and fusions

To test if the chromosome sizes tend to have more balanced chromosome sizes, we calculated the coefficient of variation of chromosome length (CVCL) that underwent fusion and fission events and compared it with the CVCL distribution of random simulations. CVCL = SD of Chromosome Length / Mean Chromosome Length. The requirements of fission simulation are as follow: 1) Break the original chromosomes to the number of target chromosomes; 2) Allow fusion events with 20% probability, i.e., times of fusion events in one simulation follows a binomial probability function with *n*= Target chromosome number - Original chromosomes number - 1 and *p*=0.2; 3) Then the number of fission events should be Target chromosome number - Original chromosomes number + Number of fusion events; 4) The smallest fragment cannot be smaller than 1Mbp after breaking; 5) The new chromosomes have the same genome size as the target chromosomes. The requirements of fusion simulation are as follows: 1) Each original chromosome appears in the target genome at least once; 2) Choose the original chromosomes randomly to fuse into the target chromosome number; 3) Fissions are allowed in some cases with 0.2 probability, specifically *R. radicans* → *R. ridleyi*; 4) The new chromosomes have the same genome size as the target chromosomes. 100,000 simulations were conducted for each pair of species and CVCL was calculated for each simulation. Then we calculated the Z-score of the observed CVCL and the CVCL distribution of 100,000 simulations. The simulation script can be found in our project GitHub page: https://github.com/Raina-M/HoloRecomb_project.

### Single-pollen nuclei library preparation

For pollen-based CO detection (**Supplementary Fig. 4a**), pollen grains were released by mechanical disruption from mature flowers or anthers of the different *Rhynchospora* species and collected into Woody Pollen Buffer (scWPB: 200mM Tris-HCl; 4mM MgCl_2_; 2mM EDTA; 86mM NaCl; 10mM Na_2_S_2_O_5_; 250mM Sucrose; 0.5mM Spermine; 0.5mM Spermidine; 1% PVP-10). To remove large debris, the obtained solution was filtered through a 50µm CellTrics cell strainer. The pollen sample was then pelleted by centrifugation and stored at −70°C until further use.

The frozen pollen samples were thawed on ice and collected into a 5µm CellTrics cell strainer. Pollen grains were macerated inside the cell strainer, releasing the pollen nuclei. The Pollen nuclei were recovered from the macerated pollen by adding scWPB with some additives (scWPC; 5mM DTT; 1%BSA; 0.2 U/µL Protector RNase Inhibitor). The pollen nuclei solution was stained with DAPI (1µg/µl) and used for Fluorescence-activated cell sorting (FACS) by employing BD FACSAriaIII Fusion Flow Cytometer with a 70µm nozzle. Nuclei were dispensed into a 96-well plate containing Collection Buffer (1xPBS, 1%BSA, and 0.2 U/µL Protector RNase Inhibitor). The quality and number of nuclei were evaluated with the LUNA-FX7 Automated Cell Counter. Nuclei solutions were used for library preparation using Chromium Next GEM Single Cell ATAC or Chromium Next GEM Single Cell 5’ Kits following the manufacturer’s instructions. Libraries sequenced at BGI Genomics following Chromium 10x Kit recommendations.

### Single-cell analysis

COs were detected using a similar pipeline reported in Catellani and Zhang, et al.^14^. An updated pipeline description is available on the GitHub page: https://github.com/Raina-M/detectCO_by_scRNAseq.

### Genome-wide mapping of CO in R. pubera

For detecting CO formation in *Rhychospora pubera* (**Supplementary Fig. 4b**), reciprocal crossings of two accessions containing 2*n* = 10 and 2*n* = 12 chromosomes were generated. Successful crosses were identified through cytological analysis by finding F1 individuals with 2*n* = 11 (5+6). Five F1 plants containing 2*n* = 11 chromosomes were identified. These F1 individuals were allowed to self-pollinate, generating an F2 segregating population. Genomic DNA from a total of 489 F2 individual plants and their respective F1 parents was extracted and deep-sequenced with an Illumina HiSeq 3000 in 150-bp paired-end mode. Alternatively, DNBseq short-read sequencing (BGI Genomics, Hong Kong) was employed. The sequencing results were analysed using our pipeline at GitHub page: https://github.com/Raina-M/HoloRecomb_project, which enabled us to map and quantify meiotic recombination events.

### Crossover interference analysis

CO interference was computed with MADpattern (v.1.1)^68^. Using an inter-interval resolution around 5Mbp, the number of inter-interval equals to the closest ceiling of the quotient of maximal chromosome size of this species divided by 5Mbp. Considering that the chromosome size of *R. cephalotes* is much smaller than in other *Rhynchospora* species, resolution inter-interval size was reduced to 3Mbp.

### Immunostaining

Immunocytochemistry experiments were carried out as in Castellani and Zhang et al.^14^, with some modifications. Briefly, young flowers of *R. pubera, R. breviuscula, R. cephalotes, R. colorata, R. ciliata* and *R. nervosa* were harvested and fixed in ice-cold 4% (w/v) paraformaldehyde (PFA) in phosphate buffered saline (PBS) solution (pH 7.5, 1.3 M NaCl, 70 mM Na2HPO4, 30 mM NaH2PO4) and 0.1% (v/v) Triton X-100 for 30 min under vacuum conditions. Flowers were dissected and anthers of the appropriate size were selected to have a high density of meiocytes in Prophase I stages, especially zygotene. Meiocytes were released from the locules by puncturing the locules with needles. This was carried out in PBS with 0.1% (v/v) Triton X-100. The suspension of meiocytes was stirred to separate individual nuclei and to remove excess debris from the slide. A coverslip was strongly pressed onto the suspension to collapse the single meiotic nuclei into a bidimensional layer, exposing epitopes. Specimens were mounted with Vectashield containing 0.2 µg/ml DAPI and checked for zygotene stages. Selected slides were incubated with a blocking buffer (3% (w/v) bovine serum albumin (BSA) in PBS + 0.1% (v/v) Triton X-100) for 1 hour at 37 °C to block and permeabilize the cells. The following antibodies were used: Anti-AtASY1 raised in rabbit (inventory code PAK006)^69^; anti-AtMLH1 raised in rabbit (inventory code PAK017)^70^; The anti-RpZYP1 was raised in rat against the peptide TAERLVKDQASVKNDLEC (Gene ID: RP1G00482580/RP4G01479980/RP2G00810370/RP5G017 38420) and affinity-purified (Lifeprotein). Primary antibodies were diluted in blocking buffer to a final dilution of 1:200. Samples were incubated with primary antibodies overnight at 4 °C. The following day, slides were washed three times for 5 min each with PBS + 0.1% (v/v) Triton X-100. Samples were incubated with secondary antibodies for 2h at room temperature. Secondary antibodies were conjugated with STAR ORANGE (Abberior, STORANGE-1007) or Alexa Fluor 488 (Thermofisher Goat anti-Rabbit IgG (H+L), Superclonal™ Recombinant Secondary Antibody, A27034) diluted 1:250 in blocking buffer. Slides were washed three times for 5 min each in PBS containing 0.1% (v/v) Triton X-100 and then allowed to dry. Slides were then re-mounted (Vectashield + 0.2 µg/ml DAPI). Meiotic cells of the desired stage were pre-imaged for subsequent FISH. Images were acquired using a Zeiss Axio Imager Z2 with an Apotome system for optical sectioning. Images were deconvolved and processed with Zen 3.2 software and Adobe Photoshop.

### Fluorescent in situ Hybridisation (FISH)

FISH was performed as described by Castellani and Zhang et al.^14^. In summary, slides of interest had their coverslips removed and were re-fixed in a solution of 4% formaldehyde in PBS and 0.1% Tween-20 for 10 min. Slides were washed 3 times for 5 min each in PBS-Triton X-100. The hybridisation mix was added to the samples before coverslips were applied and sealed with rubber cement. Denaturation was carried out at 75 °C for 10min on a hot plate. Slides were subsequently incubated overnight at 37 °C in a moist chamber. The next day, coverslips were removed, and stringency washes were performed. The sequence of washes was the following: 2XSSC+0,05% Tween-20 for 5 min at 37 °C, 20% formamide in 2XSSC+0,05% Tween-20 for 5 min at 37 °C, 2XSSC+0,05% Tween-20 for 5 min at 37 °C twice, 2XSSC+0,05% Tween-20 for 5 min at room temperature. Slides were mounted, and images were taken as described above.

### Telomere Nearest Neighbour Distance analysis

Telomere clustering was measured as follows. Images of high quality, with a clear telomeric signal and at the correct stage (early zygotene), were selected. The Telomeric signal was isolated, converted from 16-bit to 8-bit and thresholded. Nearest Neighbour Distance (NND) was calculated using the BioVoxxel^71^ suite in FIJI^72^, especially the particle distribution (2D) plugin. The measured median nearest neighbour distance was assigned to the cell, and the process was repeated for the whole dataset. A summary of the workflow is shown in **Supplementary Fig. 6a**.

### Meiotic axis length measurement

Axis length was measured in cells at the pachytene stage. Pachytene was defined as a continuous, gapless linear signal of ZYP1, detected by immunocytochemistry. Images were selected, and the ZYP1 signal was isolated, converted from 16-bit to 8-bit and thresholded in FIJI^72^. The signal was skeletonised, and the total skeleton length was measured as an estimate of the total axis length. A summary of the workflow is shown in **Supplementary Fig. 6b**.

### Hi-C analysis to estimate average DNA loop size

Hi-C reads were iteratively mapped with the GEM (version 3.6.1) mapper (77) from TADbit (version 1.0.1)^73^ using windows from 15 to 75 bp in 5-bp steps. Possible artifacts including “self-circle,” “dangling-end,” “error,” “extra dangling-end,” “too short,” “too large,” “duplicated,” and “random breaks” were filtered out. Binning was then conducted using an in-house script that imports the “HiC_data” module of TADbit to bin the mapped reads into square matrices of 5 kbp and 50 kbp resolution. All matrices were corrected and normalized to a total of 100 million interaction counts by scaling the sum of all interactions within the matrix.

HiCExplorer (version 3.7)^74^ was used to plot normalized matrices at a 50 kbp resolution and to obtain genome-wide contact probability P(s) curves at a 5 kbp resolution. Slopes were then calculated as the derivative of the contact probability versus genomic distance curves, with the highest point of the slope corresponding to the average chromatin loop size^34,75^.

### Contact domain size analysis

Deep sequencing Hi-C data were aligned to reference genome and.hic file was generated by juicer (v2.0)^76^. Hi-C contacts were extracted with juicer_tools.jar (v2.20.00) based on inter_30.hic output from juicer. The resolution of Hi-C contacts is 5kbp with VC_SQRT normalization method: “java -Xmx16g -jar juicer_tools jar dump observed VC_SQRT inter_30.hic chrID chrID BP 5000 hic_contatc.txt”. The scripts used to calculate Hi-C contact probability P(s) and P(s) slopes can be found in our project GitHub page: https://github.com/Raina-M/HoloRecomb_project. The first turning point of P(s) slope was defined as the contact domain size.

### Measurement of simulated axis length

The simulations described in the companion paper^7^ were used here to compare the axis length of meiotic chromosomes of 6 *Rhynchospora* species. We calculated the axis length as the sum of the distances between neighbor centromeric arrays, that is, between the centers of centromeric arrays *i* and *i+1*. The python scripts used to perform this analysis can be found in https://github.com/InSilicoGenebankProteomics/Rhynchospora_length.

## Data availability

Genome assemblies and raw sequencing data have been deposited in the European Nucleotide Archive under accession PRJEB103769. The reference genomes, sequencing data, annotations and all tracks presented in this work are made available for download at EDMOND, the Open Research Data Repository of the Max Planck Society. All scripts used in the present study are available at https://github.com/Raina-M/HoloRecomb_project and https://github.com/InSilicoGenebankProteomics/Rhynchospora_length. All other data are available from the corresponding author upon request.

## Acknowledgements

This study was funded by the German Research Foundation (DFG MA 9363/2-1 and MA 9363/3-1), the Max Planck Society (core funding to A.M.), and the European Union (European Research Council Starting Grant, HoloRECOMB, grant no. 101114879 to A.M.). The DFG also funded this work under Germany’s Excellence Strategy—EXC 493 2048/1–390686111 (to A.M.). M.Z. and A.S.C. are financially supported by the DFG (grant no. MA 9363/2-1 and SO 2132/1-1, respectively). This study was also funded by CONARE/Max Planck Society under the project “CENTROMERE CHARACTERIZATION OF HOLOCENTRIC RHYNCHOSPORA SPECIES” (111C3508) to granted to A.A.G and A.M. A.R.-H. was supported by the Spanish Ministry of Science and Innovation (PID2020-112557GB-I00 and PID2024-156042NB-I00), the Agència de Gestió d’Ajuts Universitaris i de Recerca, AGAUR (2021SGR00122 to ARH). L.M.-G. was supported by an FPU predoctoral fellowship from the Ministry of Science, Innovation and University (FPU18/03867). We thank Dorota Paczesniak for expert assistance with figure design.

## Author contributions

A.M. conceived the study. A.M. and M.Z. performed genome assembly and bioinformatic analyses. M.Z. performed all recombination analyses. S.S. performed all the single-cell pollen sequencing experiments. M.C. performed the cytological analysis of meiosis. M.Z., L.M.-G., A.R.-H. performed Hi-C analysis. M.Z. and J.T.L. performed synteny analysis. M.Z., L.A.R., L.M-P. and A.L.L.V. performed repeat annotation. N.G. and B.H. performed Hi-C libraries. M.M. and B.H. performed single-cell libraries. E.C., L.P.F., A.A.G., A.P-H. provided the sample material. W.W.T. taxonomically identified all plant samples. A.S.C. performed modelling analysis of axis and loop architecture. M.Z., S.S., M.C. and A.M. wrote the manuscript with the input from all other authors. All authors approved the submitted version of the manuscript.

---

**Extended Data Table 1.** 20 *Rhynchospora* species chromosome number, assembly type, genome size and sample origin. If only the haploid genome size is shown, the corresponding species usually has a highly homozygous genome, making the phasing impossible. Genome size here includes only pseudo-chromosomes.

| Species | Karyotype | Assembly type | Genome size (bp) | Plant sample source |
| --- | --- | --- | --- | --- |
| *R. rugosa* | $2n=2x=36$ | Haploid | 235,231,465 | This study |
| *R. alba* | $2n=2x=26$ | Haploid | 235,949,887 | Mata-Sucre et al.<sup>11</sup> |
| *R. cephalotes* | $2n=2x=18$ | Haplotype 1 | 303,606,085 | Mata-Sucre et al.<sup>11</sup> |
|  |  | Haplotype 2 | 303,631,510 |  |
| *R. ridleyi* | $2n=2x=12$ | Haplotype 1 | 423,345,332 | Mata-Sucre et al.<sup>11</sup> |
|  |  | Haplotype 2 | 426,461,118 |  |
| *R. gaudichaudii* | $2n=3x=15$ | Haplotype 1 | 339,693,885 | This study |
|  |  | Haplotype 2 | 343,351,133 |  |
|  |  | Haplotype 3 | 351,897,537 |  |
| *R. watsonii* | $2n=2x=10$ | Haploid | 357,361,827 | This study |
| *R. radicans* | $2n=2x=10$ | Haploid | 350,330,963 | Mata-Sucre et al.<sup>11</sup> |
| *R. pubera* | $2n=2x=10$ | Haploid | 1,694,878,336 | Hofstatter et al.<sup>6</sup> |
| *R. breviuscula* | $2n=2x=10$ | Haplotype 1 | 368,174,147 | Hofstatter et al.<sup>6</sup> |
|  |  | Haplotype 2 | 370,478,156 |  |
| *R. nervosa* | $2n=2x=10$ | Haplotype 1 | 450,937,689 | Mata-Sucre et al.<sup>11</sup> |
|  |  | Haplotype 2 | 455,796,437 |  |
| *R. ciliata* | $2n=2x=10$ | Haplotype 1 | 462,117,430 | Mata-Sucre et |
|  |  | Haplotype 2 | 441,783,790 | al.<sup>11</sup> |
| *R. colorata* | $2n=2x=10$ | Haplotype 1 | 432,477,466 | Mata-Sucre et al.<sup>11</sup> |
|  |  | Haplotype 2 | 430,351,217 |  |
| *R. tenerrima* | $2n=4x=20$ | Subgenome 1 | 232,714,659 | Mata-Sucre et al.<sup>11</sup> |
|  |  | Subgenome 2 | 242,448,059 |  |
| *R. filiformis* | $2n=2x=10$ | Haplotype 1 | 239,492,786 | Mata-Sucre et al.<sup>11</sup> |
|  |  | Haplotype 2 | 233,767,920 |  |
| *R. austrobrasiliensis* * | $2n=6x=18$ | Pseudohaplotype 1 | 347,896,131 | Zhang et al.<sup>17</sup> |
|  |  | Pseudohaplotype 2 | 402,874,894 |  |
|  |  | Pseudohaplotype 3 | 359,096,123 |  |
|  |  | Pseudohaplotype 4 | 352,179,589 |  |
|  |  | Pseudohaplotype 5 | 354,838,715 |  |
|  |  | Pseudohaplotype 6 | 348,573,314 |  |
| *R. tenuis* (REC) | $2n=2x=4$ | Haplotype 1 | 348,158,620 | Zhang et al.<sup>17</sup> |
|  |  | Haplotype 2 | 354,512,391 |  |
| *R. tenuis* (JGV-16) | $2n=2x=4$ | Haplotype 1 | 377,634,235 | Zhang et al.<sup>17</sup> |
|  |  | Haplotype 2 | 349,658,151 |  |
| *R. tenuis* (JGV-17) | $2n=2x=4$ | Haplotype 1 | 377,385,800 | Zhang et al.<sup>17</sup> |
|  |  | Haplotype 2 | 348,979,334 |  |
| *R. tenuis* (JGV-89) | $2n=2x=4$ | Haplotype 1 | 377,247,412 | Zhang et al.<sup>17</sup> |
|  |  | Haplotype 2 | 349,182,916 |  |
| *R. tenuis* (PECP-47) | $2n=2x=4$ | Haplotype 1 | 381,933,207 | Zhang et al.<sup>17</sup> |
|  |  | Haplotype 2 | 349,003,858 |  |
| *R. tenuis* (PECP-48) | $2n=2x=4$ | Haplotype 1 | 382,648,587 | Zhang et al.<sup>17</sup> |
|  |  | Haplotype 2 | 350,935,085 |  |
| *R. tenuis* (JGV-35-6) | $2n=2x=4$ | Haplotype 1 | 382,455,145 | Zhang et al.<sup>17</sup> |
|  |  | Haplotype 2 | 350,070,891 |  |
| *R. tenuis* (JGV-36) | $2n=2x=4$ | Haplotype 1 | 382,926,756 | Zhang et al.<sup>17</sup> |
|  |  | Haplotype 2 | 348,977,203 |  |
| *R. tenuis* (JGV-36-7) | $2n=2x=4$ | Haplotype 1 | 383,444,050 | Zhang et al.<sup>17</sup> |
|  |  | Haplotype 2 | 349,631,601 |  |
| *R. riparia* | $2n=2x=10$ | Haplotype 1 | 297,062,901 | Mata-Sucre et al.<sup>11</sup> |
|  |  | Haplotype 2 | 286,276,137 |  |
| *R. barbata* | $2n=2x=10$ | Haplotype 1 | 514,099,728 | Mata-Sucre et al.<sup>11</sup> |
|  |  | Haplotype 2 | 514,099,728 |  |
| *R. holoschoenoides* | $2n=2x=10$ | Haplotype 1 | 254,695,320 | Mata-Sucre et al.<sup>11</sup> |
|  |  | Haplotype 2 | 255,930,137 |  |
| *R. corymbosa* | $2n=2x=18$ | Haplotype 1 | 348,126,674 | This study |
|  |  | Haplotype 2 | 338,896,338 |  |

\**R. austrobrasiliensis* has an autohexaploid genome, so six haplotypes are barely distinguishable solely based on the genomic method. The pseudo-haplotype shown here is based on the *k*-mer distance clustering from Zhang et al.<sup>17</sup>.

**Extended Data Table 2:** Karyotype changes in 20 *Rhyncospora* species and break point sources. The paired original and derived species do not indicate the real evolutionary event. The one closest to the ancestral karyotype is regarded as the original karyotype. Detailed rearrangement junction features are shown in **Supplementary Dataset 2**.

| Original | Derived | Fission | Fusion | Total |
| --- | --- | --- | --- | --- |
| *R. alba* | *R. rugosa* | 12 | 7 | 19 |
| *R. cephalotes* | *R. alba* | 13 | 7 | 20 |
| *R. gaudichaudii* | *R. cephalotes* | 5 | 1 | 6 |
| *R. radicans* | *R. pubera* | 0 | 15* | 15 |
| *R. filiformis* | *R. austrobrasiliensis* | 0 | 2 | 2 |
| *R. austrobrasiliensis* | *R. tenuis* | 0 | 1 | 1 |
| *R. holoschoenoides* | *R. corymbosa* | 5 | 1 | 6 |
| Total |  | 35 | 34 | 69 |

\*Based on Hofstatter et al. 2022<sup>6</sup>, the 15 end-to-end fusions is hypothetically from 7 independent fusion events by tracing the karyotype-evolving history.

**Extended Data Table 3:** Detailed information for the recombination data generated in this study.

| Species | Reference haplotype | Hap1 genome size | Hap2 genome size | Single cell library type | Libray ID | Cell count | CO count | Mean CO number/cell | Mean CO rate | Cell count* | CO count* | Mean CO number/cell * | CO rate * |
| --- | --- | --- | --- | --- | --- | --- | --- | --- | --- | --- | --- | --- | --- |
| *R. barbata* | haplotype 1 | 518246442 | 518350772 | scATAC | 6160.B | 2495 | 6671 | 2.673747495 | 0.515922017 |  |  |  |  |
| *R. breviuscula* | haplotype 1 | 368174147 | 370478156 | scRNA 3' | 4984.A | 1641 | 4047 | 2.466179159 | 0.669840395 |  |  |  |  |
| *R. cephalotes* | haplotype 1 | 303606085 | 303631510 | scRNA 3' | 5419.A** | 325 | 1439 | 4.427692308 | 1.458367446 | 2719 | 12016 | 4.419271791 | 1.455594 |
|  |  |  |  | scATAC | 5999.D | 58 | 273 | 4.706896552 | 1.55033011 |  |  |  |  |
|  |  |  |  | scATAC | 6160.A | 2336 | 10304 | 4.410958904 | 1.452855895 |  |  |  |  |
| *R. ciliata* | haplotype 1 | 462117430 | 441783790 | scATAC | 5999.A | 597 | 1553 | 2.601340034 | 0.562917532 |  |  |  |  |
| *R. colorata* | haplotype 1 | 432477466 | 430351217 | scRNA 3' | 5491.A** | 429 | 1004 | 2.34032634 | 0.541144111 |  |  |  |  |
| *R. nervosa* | haplotype 2 | 450937689 | 455796437 | scRNA 5' | 6385.A | 1173 | 2984 | 2.543904518 | 0.55812295 |  |  |  |  |
| *R. pubera* *** | haplotype 1 | 1472673118 |  | WGS F2 |  | 489 | 1158 |  |  |  |  |  |  |

\*Summary of all libraries of this species.
\*\*Library 5491.A appear twice (*R. cephalotes* and *R. colorata*) because we mixed the pollens from two species for multiplexing, the species identity of single cells were assigned with bioinformatic method (Castellani and Zhang<sup>14</sup>).
\*\*\**R. pubera* CO detection was based on F2 whole-genome DNA sequencing, so the 'cell count' here refers to the number of F2 individual plants.

## Extended Data Figures

**Extended Data Fig. 1:**
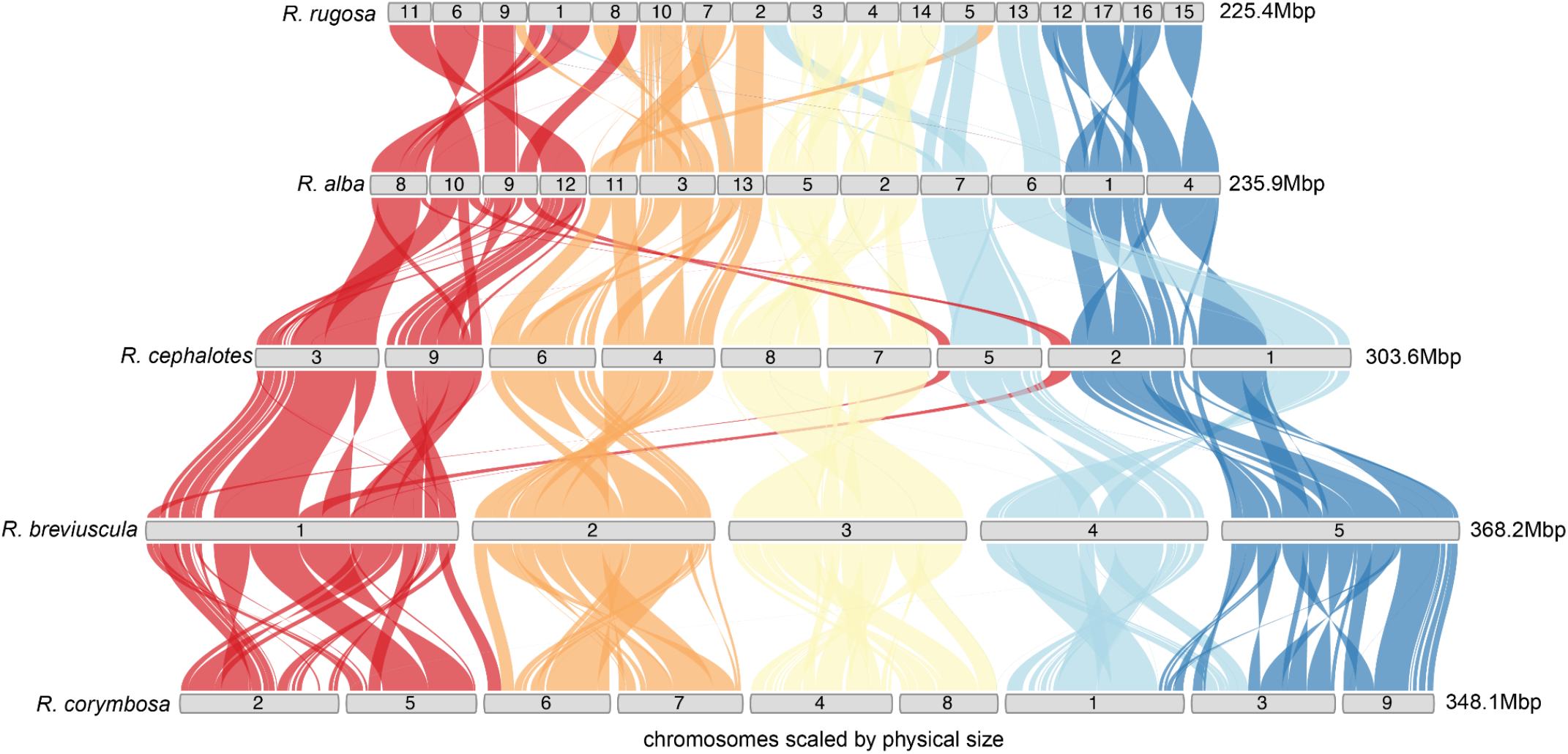
Agmatoploid karyotype evolution in *Rhynchospora*. DEEPSPACE synteny reconstruction of chromosome number evolution from the inferred *Rhynchospora* ancestral karyotype (RAK) with *n* = 5 (i.e., *R. breviuscula*). Recurrent chromosome fissions doubled chromosome number without genome duplication, generating an intermediate agmatoploid state with *n* = 10. Subsequent chromosome fusion events independently reduced chromosome number to *n* = 9 in *R. corymbosa* and *R. cephalotes*. Additional rounds of chromosome fission further increased chromosome number, producing the karyotype of *R. alba* (*n* = 13) and, through continued fragmentation, the highly derived karyotype of *R. rugosa* (*n* = 18), representing an approximate doubling relative to *R. cephalotes*. Throughout this process, large syntenic blocks are retained, indicating that chromosome number change predominantly reflects structural fragmentation and fusion rather than polyploidy.

**Extended Data Fig. 2:**
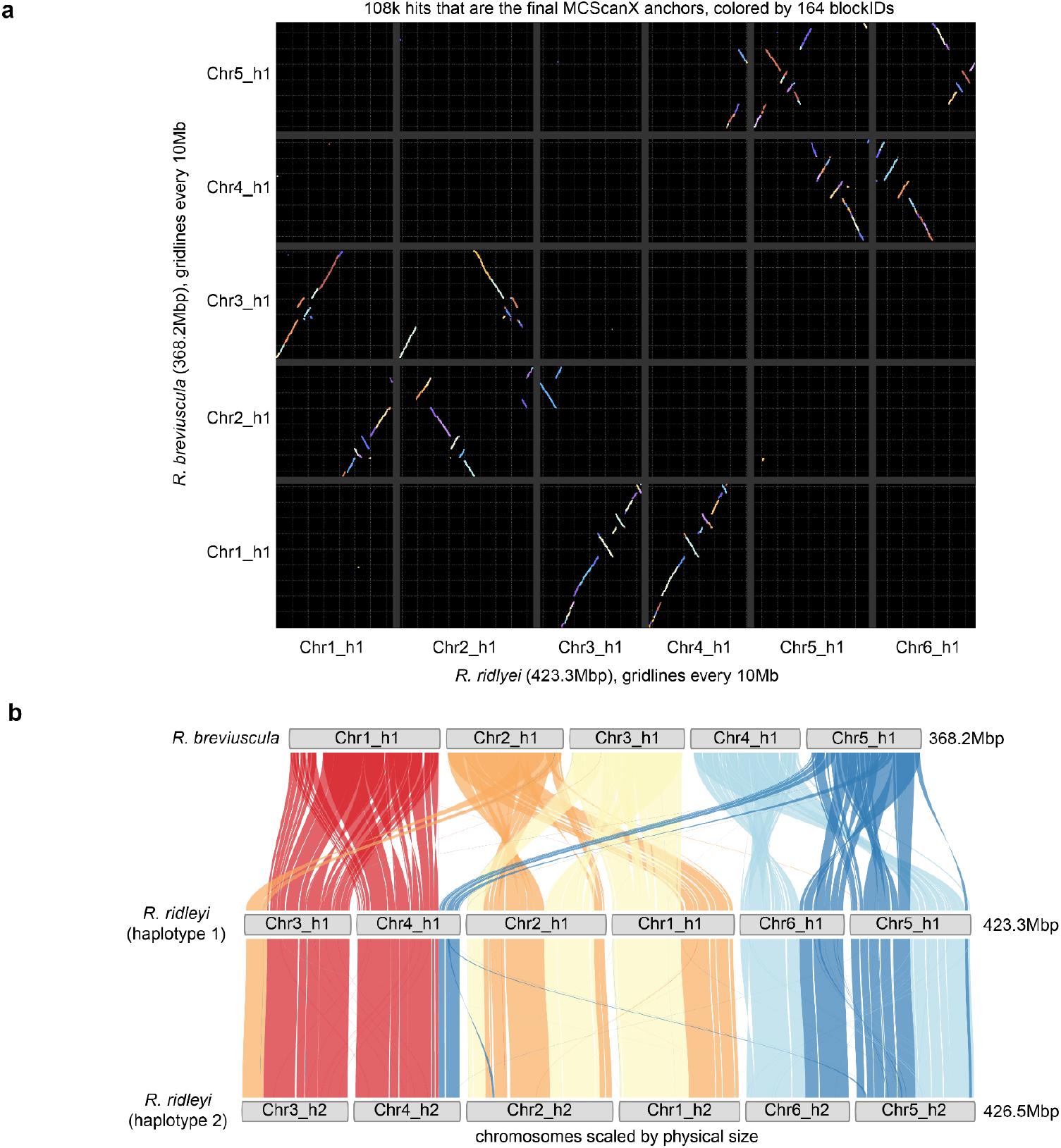
Whole-genome duplication followed by extensive chromosome rearrangement and simploidy in *Rhynchospora ridleyi*. (**a**) DEEPSPACE dot-plot representation of gene-based synteny between *R. breviuscula* (*n* = 5) and *R. ridleyi* haplotype 1 (n = 6), based on MCScanX anchors. Each dot represents a syntenic gene pair, coloured by syntenic block identity, revealing the presence of duplicated syntenic segments consistent with a recent whole-genome duplication in *R. ridleyi*, followed by extensive intrachromosomal rearrangements. (**b**) DEEPSPACE Synteny ribbon plots comparing *R. breviuscula* chromosomes with both haplotypes of *R. ridleyi*. Coloured ribbons represent conserved syntenic blocks, illustrating genome-wide duplication of ancestral chromosomes and subsequent chromosome number reduction through multiple fusions and rearrangements, resulting in a reduced karyotype despite the underlying polyploid origin. Together, these analyses demonstrate that *R. ridleyi* underwent a whole-genome duplication followed by rapid fusion-driven diploidisation (simploidy), preserving large syntenic blocks while substantially reshaping chromosome structure.

**Extended Data Fig. 3:**
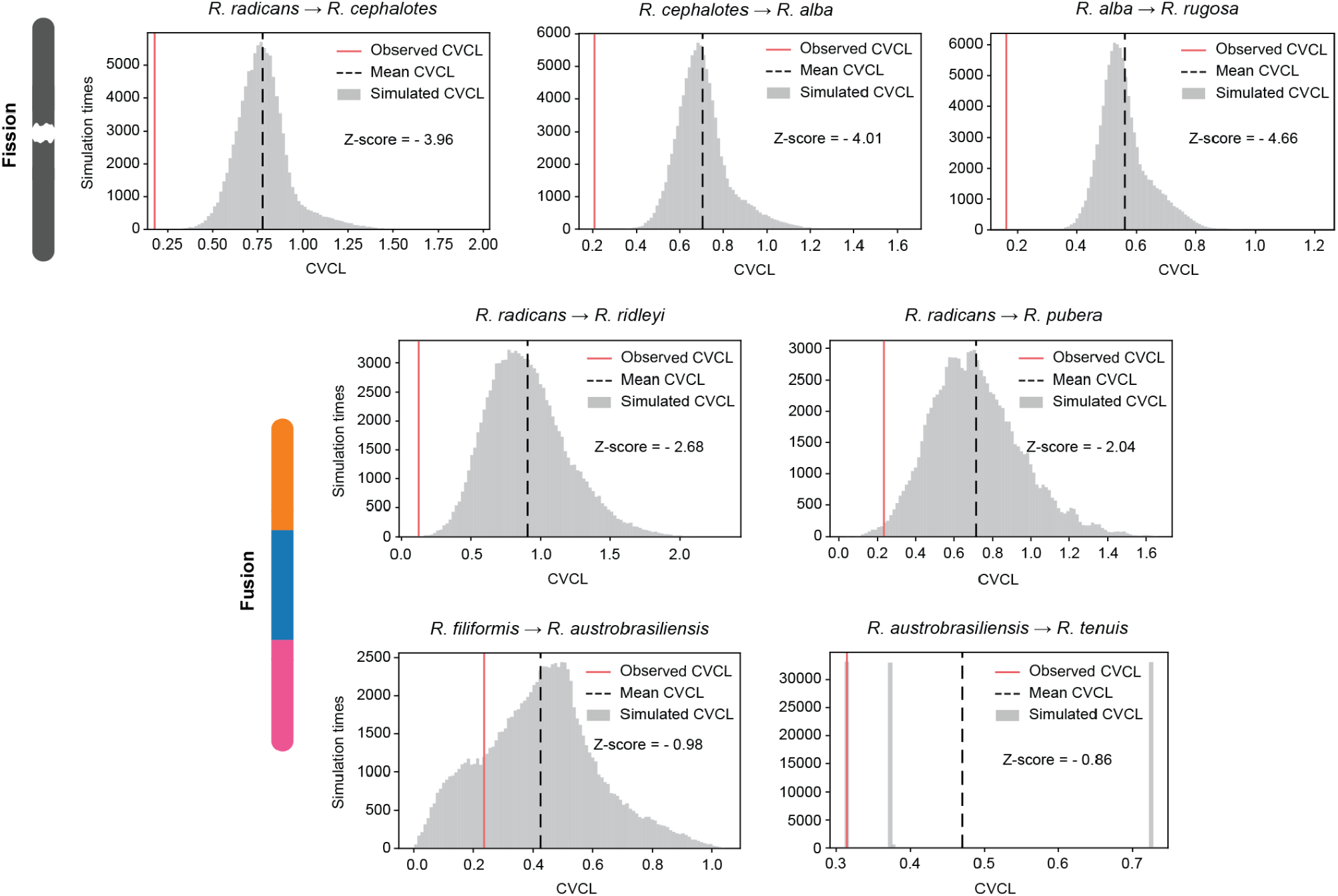
Interchromosomal size symmetry is maintained during karyotype evolution in *Rhynchospora*. Observed coefficient of variation of chromosome length (CVCL) of different *Rhynchospora* species versus the expected CVCL distribution under random chromosome-size reassortment for multiple inferred karyotype transitions across the genus (fissions: *R. radicans* → *R. cephalotes, R. cephalotes* → *R. alba, R. alba* → *R. rugosa*; fusions: *R. radicans* → *R. ridleyi, R. radicans* → *R. pubera, R. filiformis* → *R. austrobrasiliensis, R. austrobrasiliensis* → *R. tenuis*). Grey histograms show CVCL distributions obtained from the random simulation of chromosome breaks and fusions, with dashed black lines indicating the mean of simulated CVCL values. Observed CVCL values for the extant target species (e.g.: *R. cephalotes* is the target species in *R. radicans* → *R. cephalotes*) are shown as solid red lines and are consistently lower than random expectations, demonstrating that chromosome fissions and fusions generate karyotypes with unusually uniform chromosome sizes. These results indicate strong constraints favouring interchromosomal size symmetry during holocentric karyotype evolution.

**Extended Data Fig. 4:**
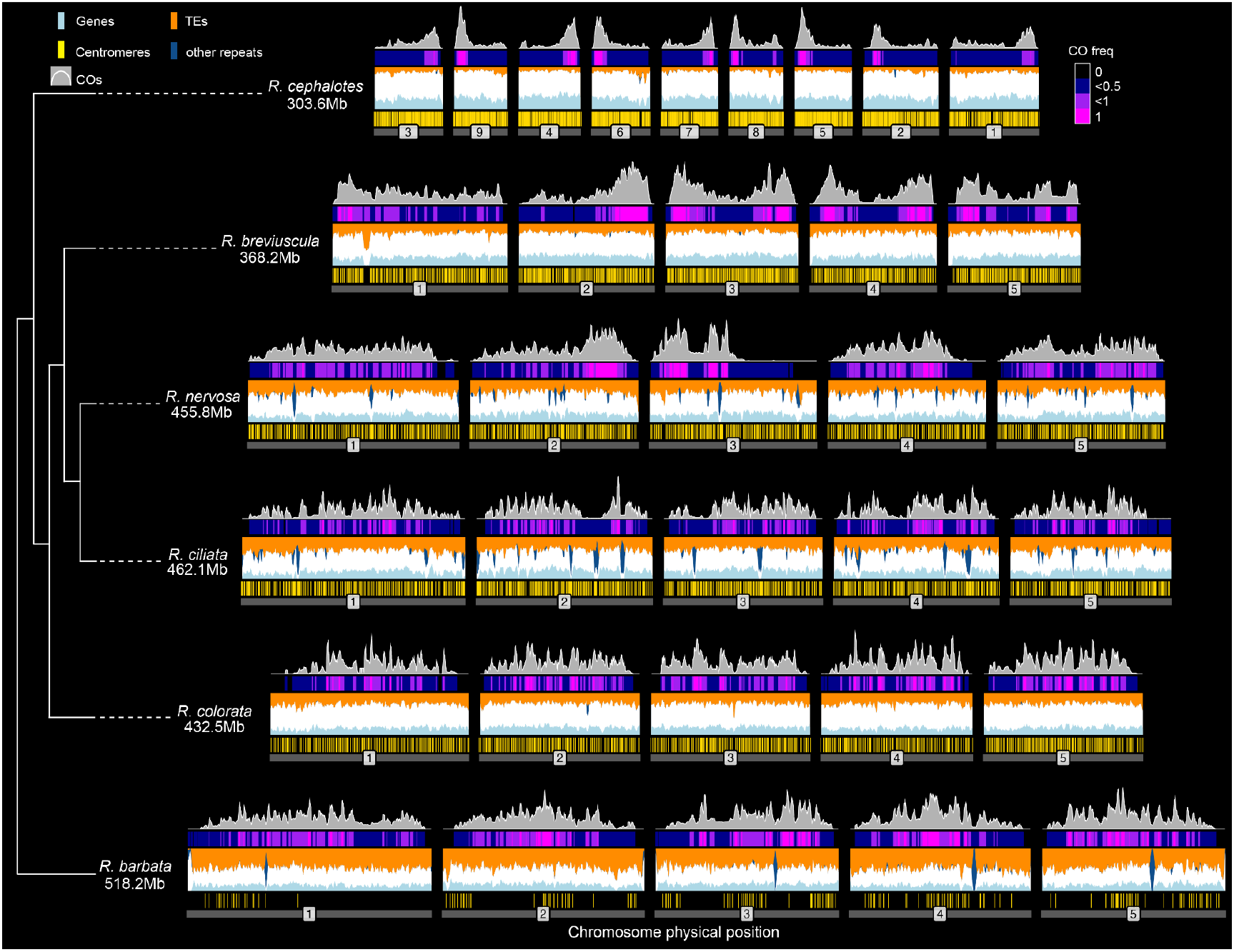
Crossover landscapes are uncoupled from local genomic features across holocentric *Rhynchospora*. Genome-wide CO frequency profiles are shown for six *Rhynchospora* species spanning distal-biased and irregular recombination landscape classes and a wide range of chromosome sizes. For each chromosome, tracks display CO distribution (grey), gene density (light blue), transposable element (TE) density (orange), centromeric *Tyba* repeat distribution (yellow), and other repetitive elements (dark blue). Despite pronounced differences in CO spatial distribution, including strong distal enrichment in some species (*R. cephalotes* and *R. breviuscula*) and irregular patterns in others (i.e., *R. nervosa, R. ciliata, R. colorata* and *R. barbata*), CO frequency shows no consistent correlation with gene density, TE content, or centromeric repeat distribution. This lack of association is observed across all chromosomes and species, regardless of recombination landscape class or chromosome size, indicating that local genomic features do not predict crossover placement in holocentric *Rhynchospora* genomes. The CO maps for *R. cephalotes, R. breviuscula, R. nervosa, R. ciliata, R. colorata*, and *R. barbata* were obtained from single-pollen nucleus sequencing. CO frequency is also shown as a heatmap above the tracks. The CO rates of each species were normalized to a 0-1 scale based on their minimum and maximum CO rates. The minimum CO rates are 0 for all species, and the maximum CO rates are 9.25, 2.34, 2.28, 2.78, 2.22, 1.94 cM/Mb, following the order of the species on the phylogenetic tree from top to bottom.

**Extended Data Fig. 5:**
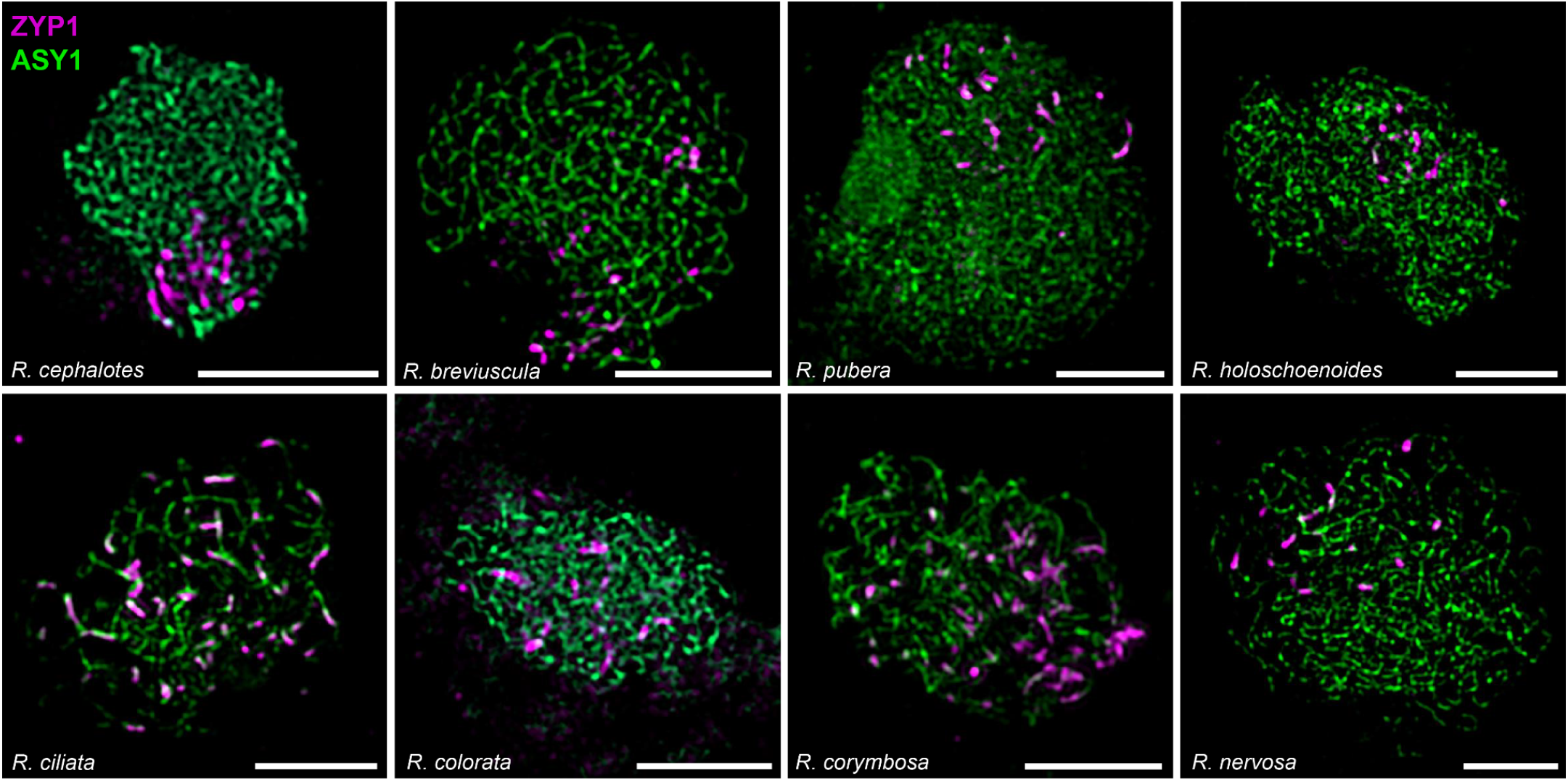
Overview of initiation of synapsis at early zygotene in all species analyzed in this study. This panel displays the behaviour of different species at the initiation of the assembly of the synaptonemal complex marked by the loading of ZYP1 (magenta) on paired chromosomes, while unpaired chromosomes are still marked by the axis component ASY1 (green). In all species ZYP1 is first loaded as short fragments. However, the distribution of these short stretches display a degree of species-specific variability that we propose correlates with different types of CO distribution. While species with distally biased crossovers (*R. pubera* n=44, *R. cephalotes* n=27, *R. breviuscula* n=16) initiate synapsis in a constrained region of their nucleus, species with irregular crossover distribution (*R. nervosa* n=27, *R. ciliata* n=21, *R. colorata* n=29) show more dispersed and less defined synapsis initiation and progression. As we do not possess recombination maps for *R. holoschoenoides* and *R. corymbosa*, based on their synapsis behaviour we would expect the first to fall in the “biased”category, and the latter to fall in the “irregular” category (respectively n=15 and n=23). Samples were prepared and images were acquired and processed as described in our methods. Scale bars correspond to 5μm.

**Extended Data Fig. 6:**
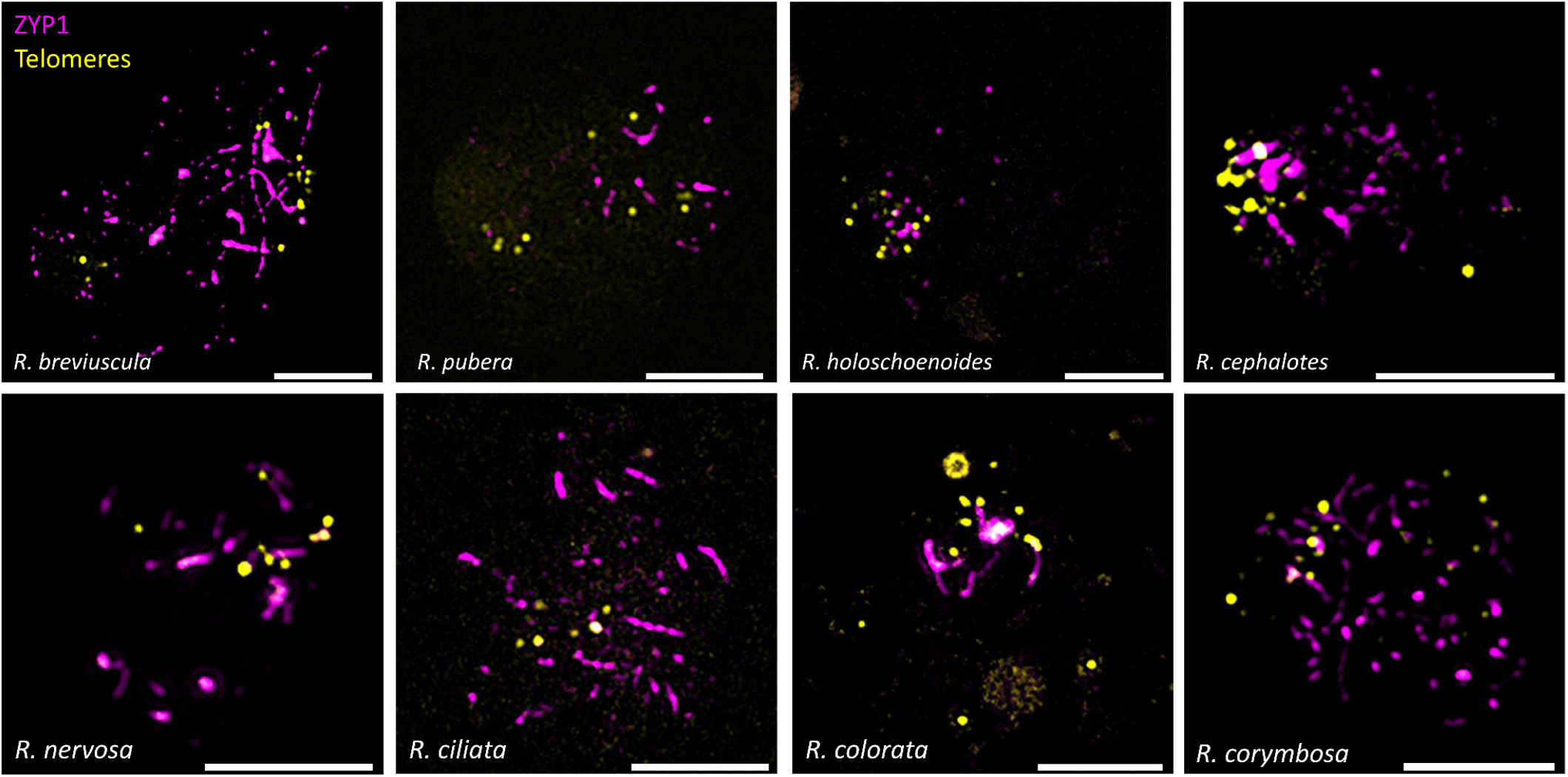
Overview of telomere positioning and synapsis initiation in all species analyzed. The figure displays the onset of synapsis related to telomere positioning during zygotene in *R. breviuscula* (n=35), *R. pubera* (n=22), *R. holoschoenoides* (n=12), *R. cephalotes* (n=7), *R. nervosa* (n=18), *R. ciliata* (n=16), *R. colorata* (n=23) and *R. corymbosa* (n=10). In all species telomeres cluster into a bouquet-like structure. A progressively loose clustering is observed in species with a biased recombination landscape, however the spatiotemporal dynamics of this process are difficult to pinpoint. We cannot rule out that subtle differences could be present between species and also as zygotene progresses, as chromosomes can greatly alter their movement. Samples were prepared and images were acquired and processed as described in our methods. Yellow signals represent telomeres, while ZYP1 (synaptonemal complex) is stained in magenta for all images. Scale bars correspond to 5μm.

**Extended Data Fig. 7:**
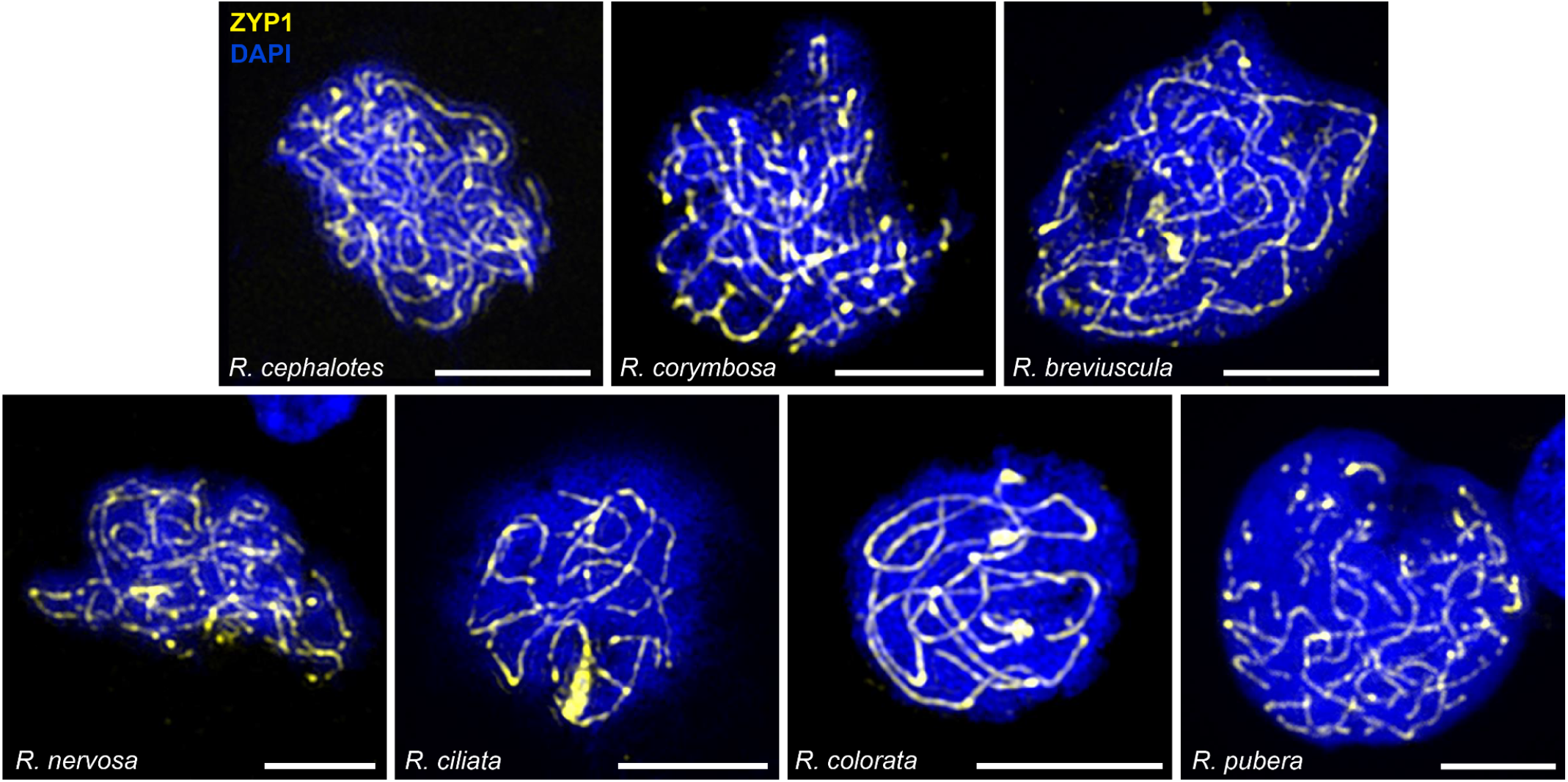
Overview of pachytene stages and ZYP1 immunostaining with all species analysed. The figure offers an overview of pachytene stages of *R. cephalotes* (n=12), *R. corymbosa* (n=13), *R. breviuscula* (n=23), *R. nervosa* (n=18), *R. ciliata* (n=13), *R. colorata* (n=12), *R. pubera* (n=10). All species are considered have similar meiosis progression and ZYP1 immunolocalisation at pachytene. Remarkably, in *R. pubera* we never found full synapsis, a feature we hypothesize might be related to an adaptation to its highly duplicated genome^6^, therefore, directly influencing its total axis length measurement. Samples were prepared and images were acquired and processed as described in our methods. Yellow signals represent ZYP1 (synaptonemal complex), while chromatin is stained with DAPI (blue) for all images. Scale bars correspond to 5μm.

## Supplementary Material

**Supplementary Fig. 1:**
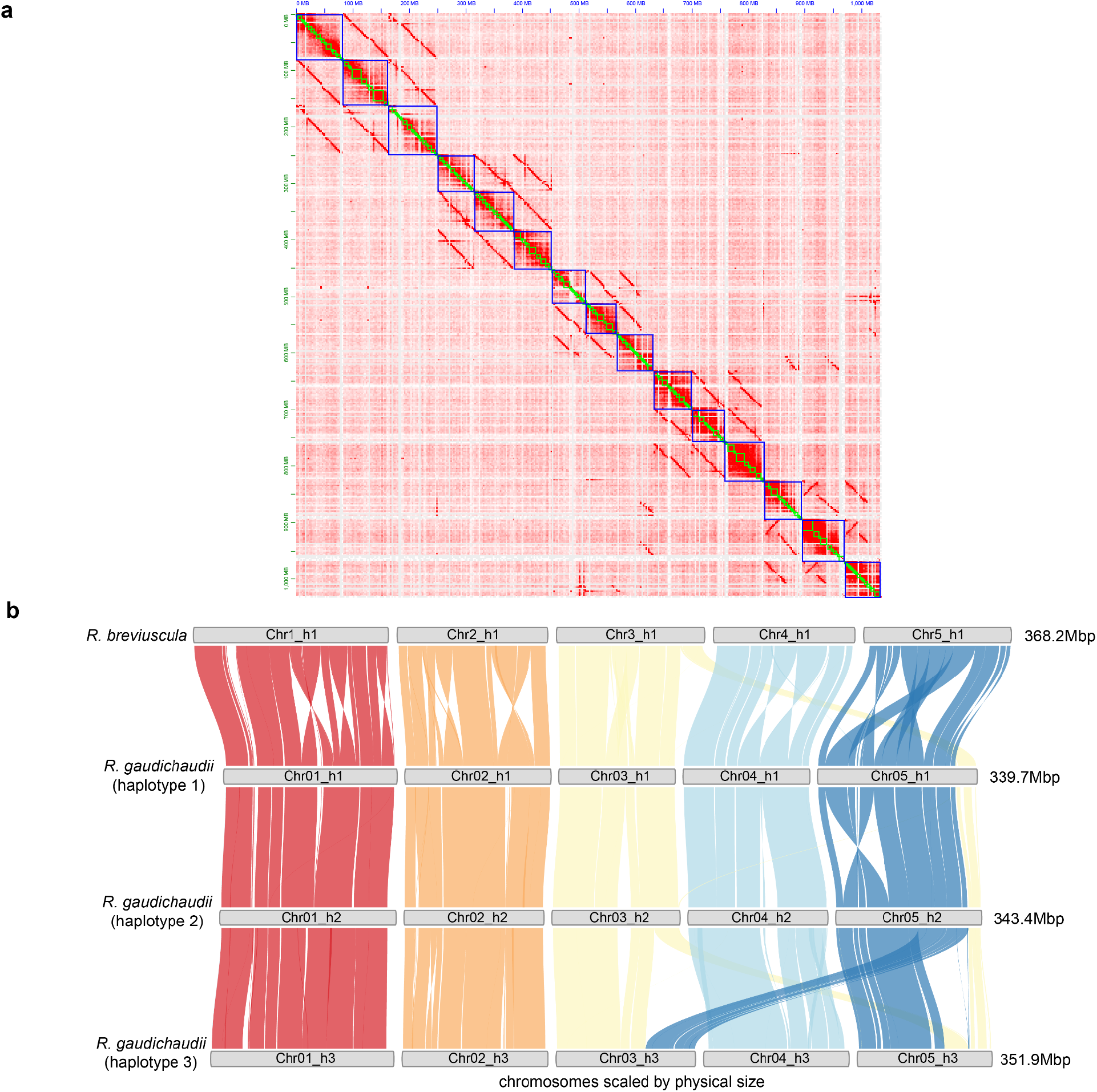
Autotriploid genome architecture of *Rhynchospora gaudichaudii*. (**a**) Whole-genome Hi-C contact map of *R. gaudichaudii* showing three phased chromosome sets resolved using CPhasing, consistent with a triploid genome. (**b**) DEEPSPACE Synteny ribbon plots comparing *R. breviuscula* (*n* = 5) with the three phased haplotypes of *R. gaudichaudii*. Coloured ribbons represent conserved syntenic blocks, revealing high collinearity across all three haplotypes and major translocations between Chr3 and Chr5. Despite extensive efforts, subgenomes could not be distinguished based on sequence divergence or structural features, supporting a likely autotriploid origin rather than allopolyploidy.

**Supplementary Fig. 2:**
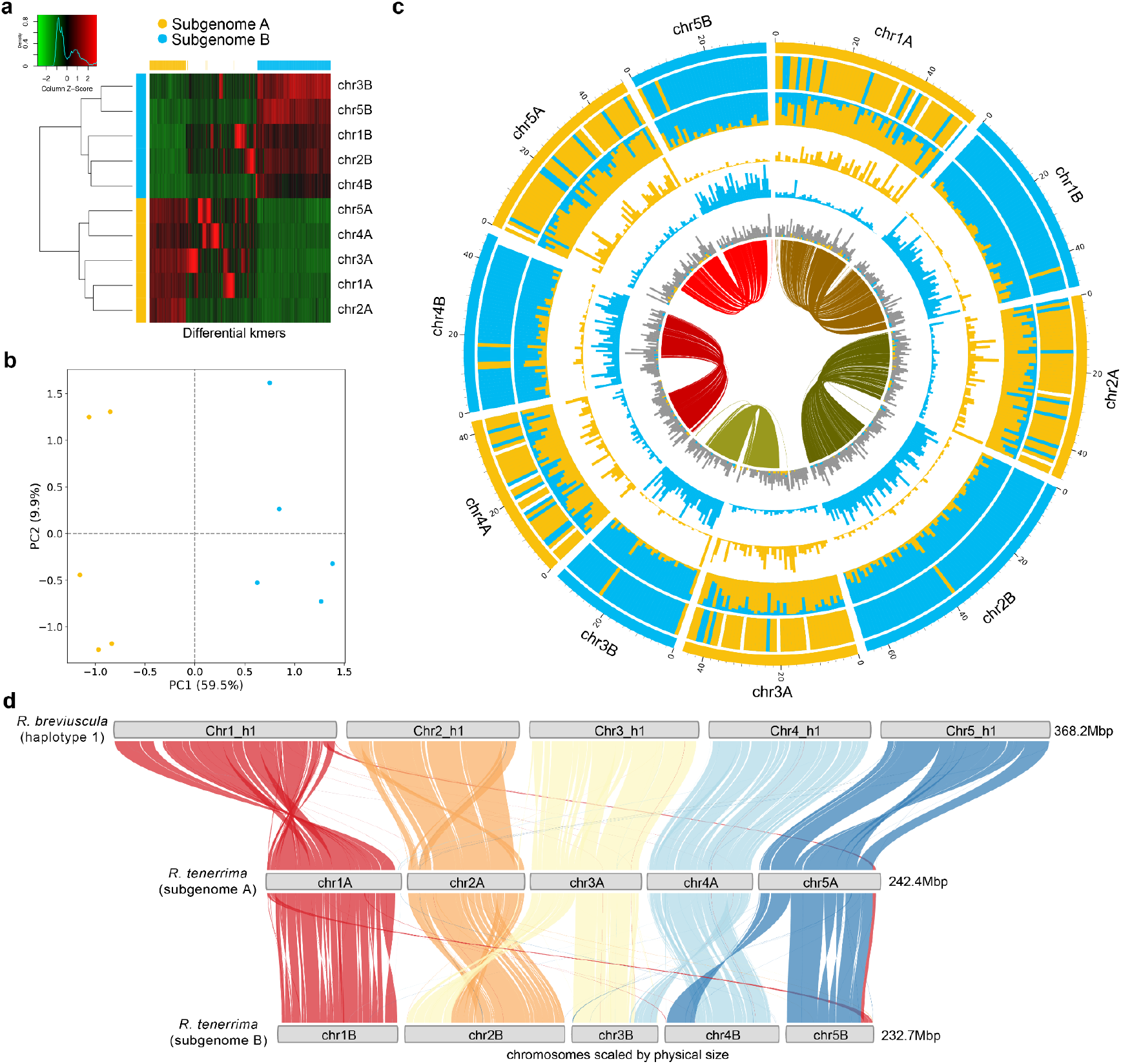
Allotetraploid genome architecture and subgenome differentiation in *Rhynchospora tenerrima*. (**a**) Heatmap of differential *k*-mer abundance across chromosomes, revealing two clearly distinct subgenomes (A and B) phased using the SubPhaser pipeline. Chromosomes cluster by subgenome identity, indicating substantial sequence divergence between the chromosomes from each subgenome. (**b**) Principal component analysis of *k*-mer frequencies further separates chromosomes into two discrete clusters corresponding to subgenomes A and B, confirming their allotetraploid origin. (**c**) Circular representation of the *R. tenerrima* genome showing chromosome organisation by subgenome. Tracks indicate subgenome assignment, *k*-mer density, and structural features along each chromosome. Inner ribbons highlight interchromosomal synteny. (**d**) DEEPSPACE Synteny ribbon plots comparing *R. breviuscula* with the two phased subgenomes of *R. tenerrima*. Conserved syntenic blocks demonstrate genome-wide duplication consistent with tetraploidy, while multiple interchromosomal rearrangements and translocations are evident, particularly within subgenome B. Together, these analyses establish *R. tenerrima* as an allotetraploid species with structurally differentiated subgenomes and asymmetric post-polyploid genome evolution.

**Supplementary Fig. 3:**
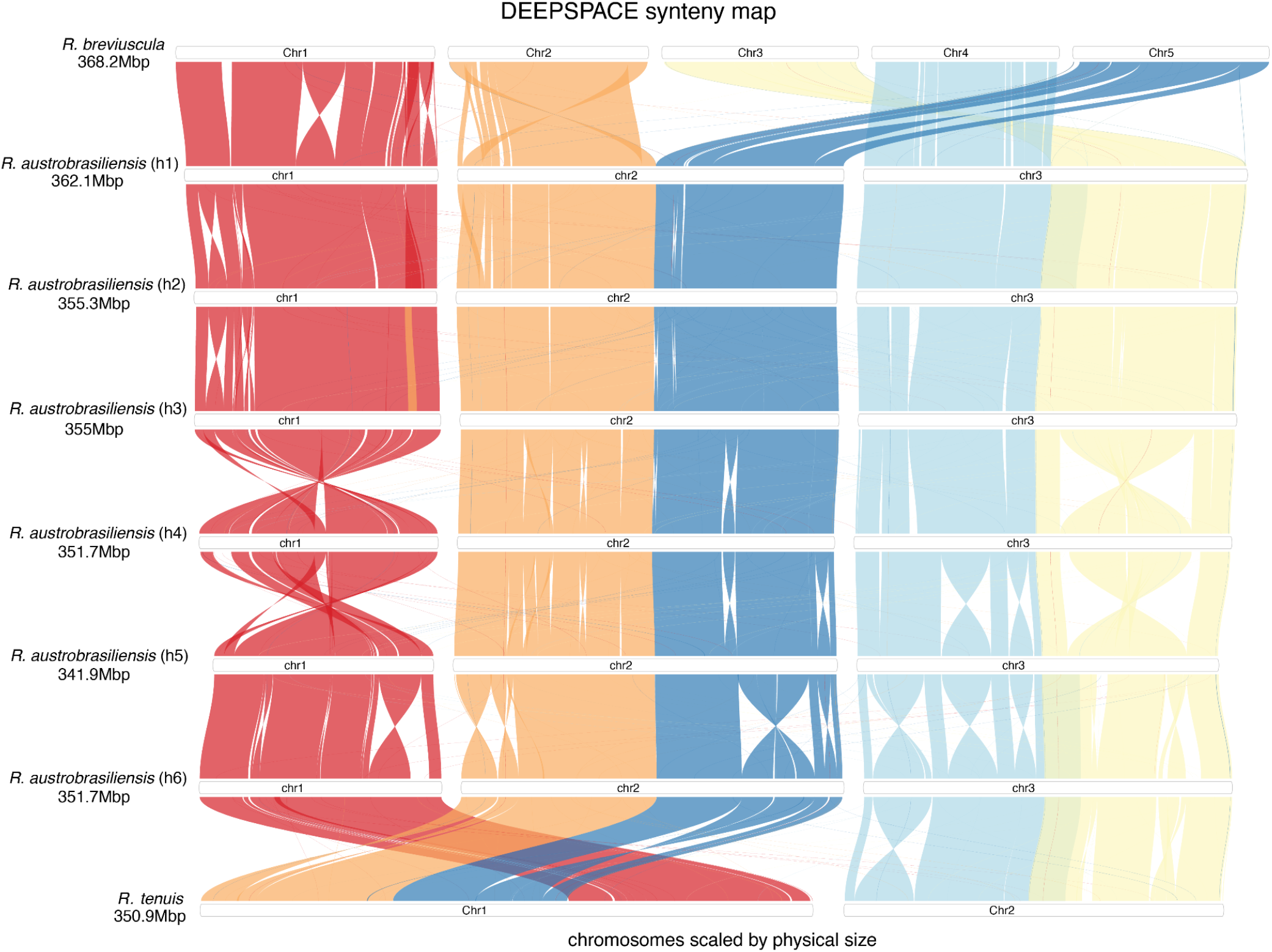
Stepwise chromosome fusion underlying karyotype reduction in *Rhynchospora austrobrasiliensis* and *R. tenuis*. DEEPSPACE synteny map comparing *R. breviuscula* (*n* = 5), multiple haplotypes of the autohexaploid *R. austrobrasiliensis*, and *R. tenuis* (*n* = 2). Chromosomes and scaffolds are scaled by physical size, and coloured ribbons indicate conserved syntenic blocks. The six phased haplotypes of *R. austrobrasiliensis* retain extensive collinearity with the ancestral *R. breviuscula* chromosomes, while showing progressive fusion of syntenic blocks. Comparison with *R. tenuis* reveals that the *R. austrobrasiliensis* karyotype preserves intermediate fusion states, supporting a stepwise chromosome fusion pathway leading to the highly reduced chromosome number of *R. tenuis*.

**Supplementary Fig. 4:**
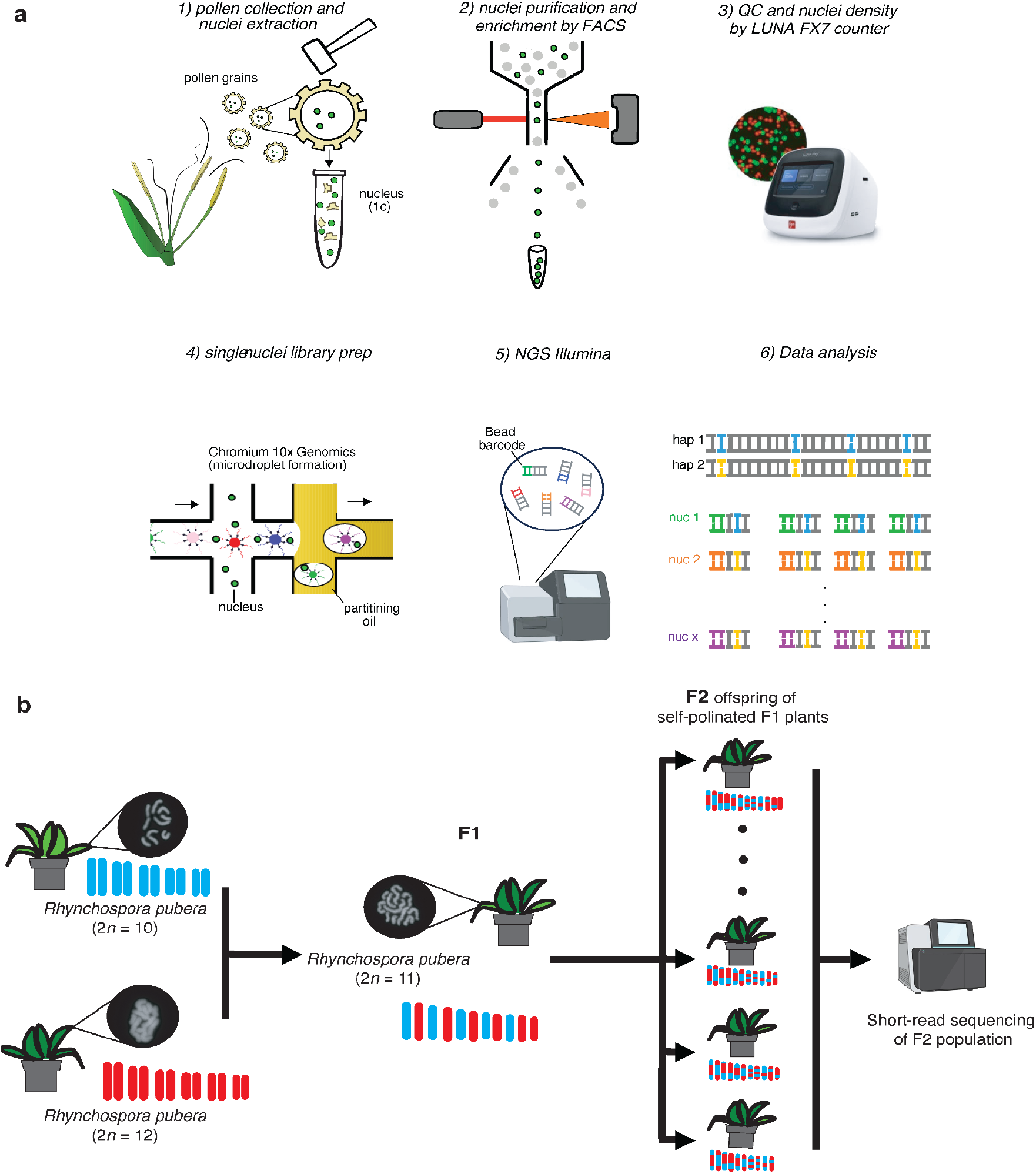
Diagram describing the CO detection pipeline in *Rhynchospora*. (**a**) Pipeline for pollen-based genotyping. To extract the pollen nuclei, pollen was macerated (1), followed by FACs enrichment of the nuclei population (2). The number and quality of nuclei were assessed using an automated counter (3) and subsequently used to prepare a single-nuclei library (4). Libraries were sequenced following kit recommendations (5) and obtained data analysed (6) according to our previously published bioinformatic pipeline^14^(**b**) Pipeline based on plant genotyping. Plants of *Rhynchospora pubera* containing either 10 chromosomes (blue) or 12 chromosomes (red) were crossed, generating a F_1_ hybrid plant with 11 chromosomes. The offspring (F_2_) of that plant were later genotyped to detect meiotic recombination events by WGS.

**Supplementary Fig. 5:**
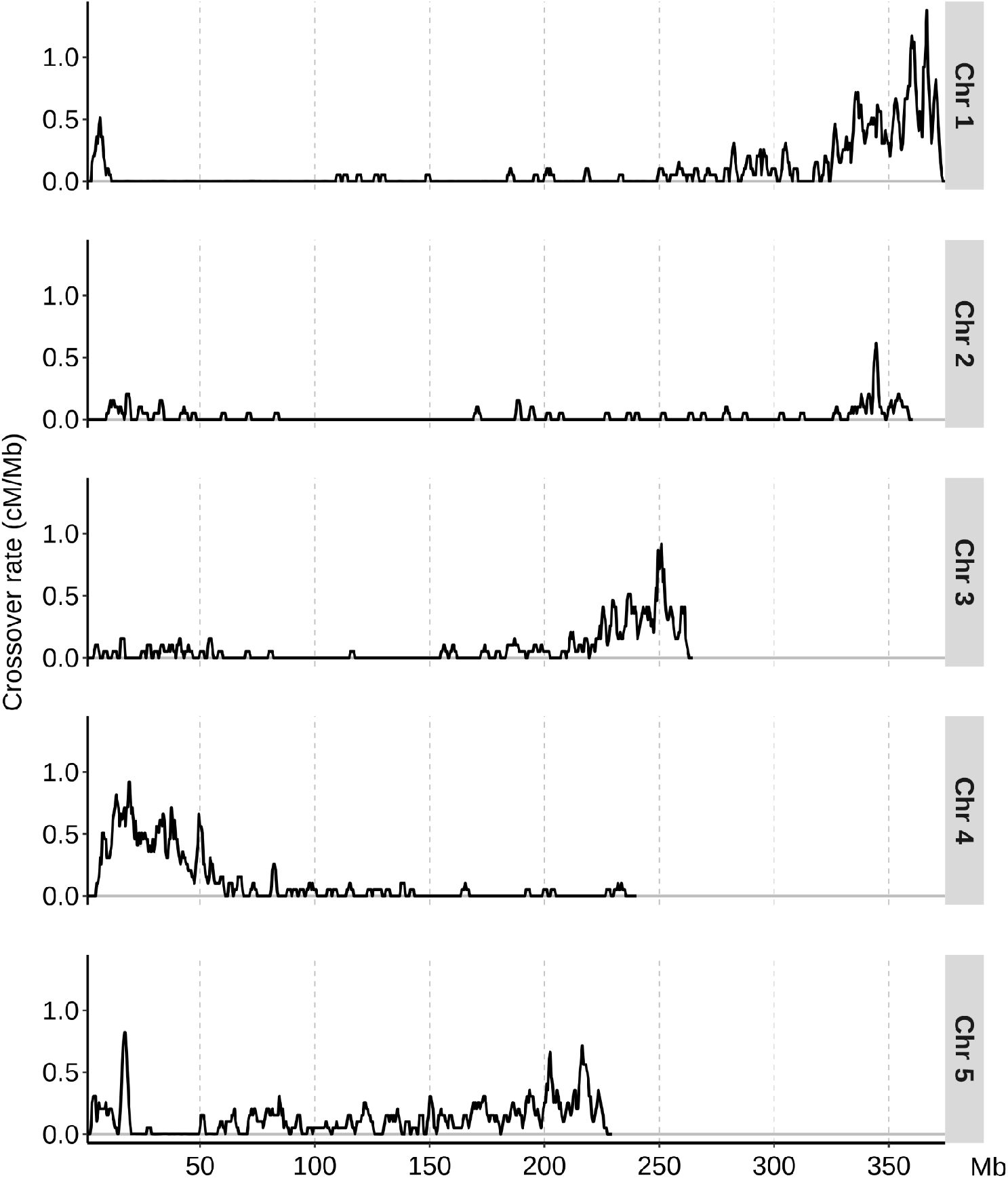
Recombination landscape of *Rhynchospora pubera* inferred from an intercytotype F_2_ mapping population. Chromosome-scale recombination landscapes (cM/Mb) for the five chromosomes of *R. pubera*, inferred from an F_2_ segregating population derived from a cross between two inbred cytotypes with different chromosome numbers (*R. pubera* red, 2*n* = 10; *R. pubera* purple, 2*n* = 12). The cross generates a fertile hybrid with 2*n* = 11, allowing genetic mapping despite karyotypic heterozygosity. While structural differences between the parental cytotypes likely contribute to variation in the absolute number of crossovers observed across chromosomes, the overall spatial distribution of recombination along chromosome arms remains consistent, with crossovers predominantly confined to distal regions. These data indicate that karyotypic rearrangements affect crossover frequency but do not substantially alter the broad-scale recombination landscape in *R. pubera*.

**Supplementary Fig. 6:**
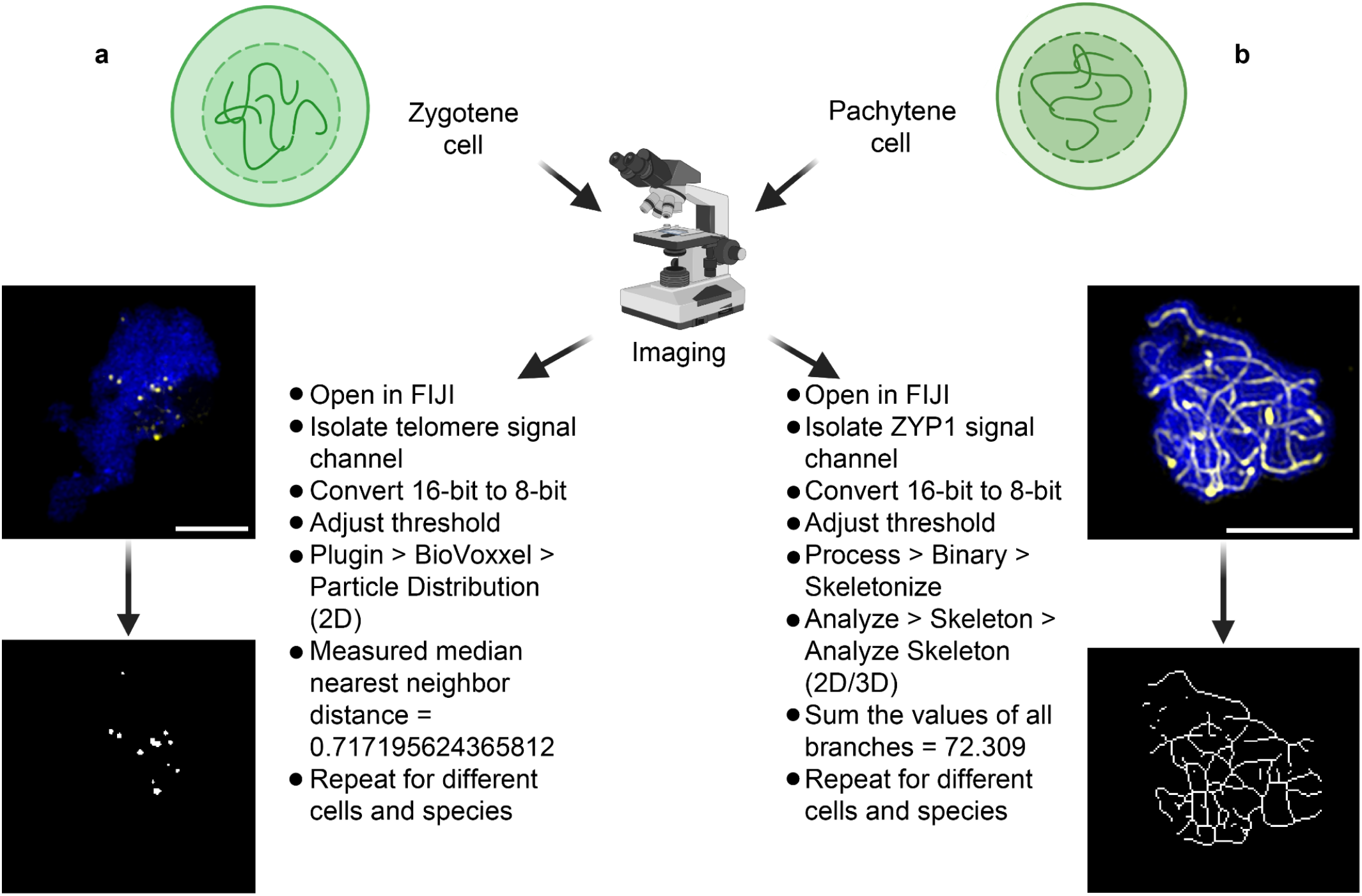
Workflow for the analysis of microscopy images for telomere nearest neighbor distance (NND) and axis length measurements. After immunocytochemistry, cells of interest with clear and explicit signals and at the correct stage were selected for image processing. Microscopy images were opened in FIJI^72^ and their channel of interest isolated, converted from 16-bit and 8-bit and thresholded. (**a**) For NND, the BioVoxxel^71^ plugin was used for 2D particle distribution analysis that returned an array of values, including the median nearest neighbour distance, which was taken as an estimation of the degree of clustering of telomeres. (**b**) For axis length, the linear signals of ZYP1 were skeletonised and the resulting skeleton was measured. The lengths of all the branches were summed and taken as an estimation of the total axis length of the cell. Yellow signal corresponds to telomeres (**a**) and ZYP1 (**b**) against DAPI (blue). Scale bars correspond to 5μm.

**Supplementary Fig. 7:**
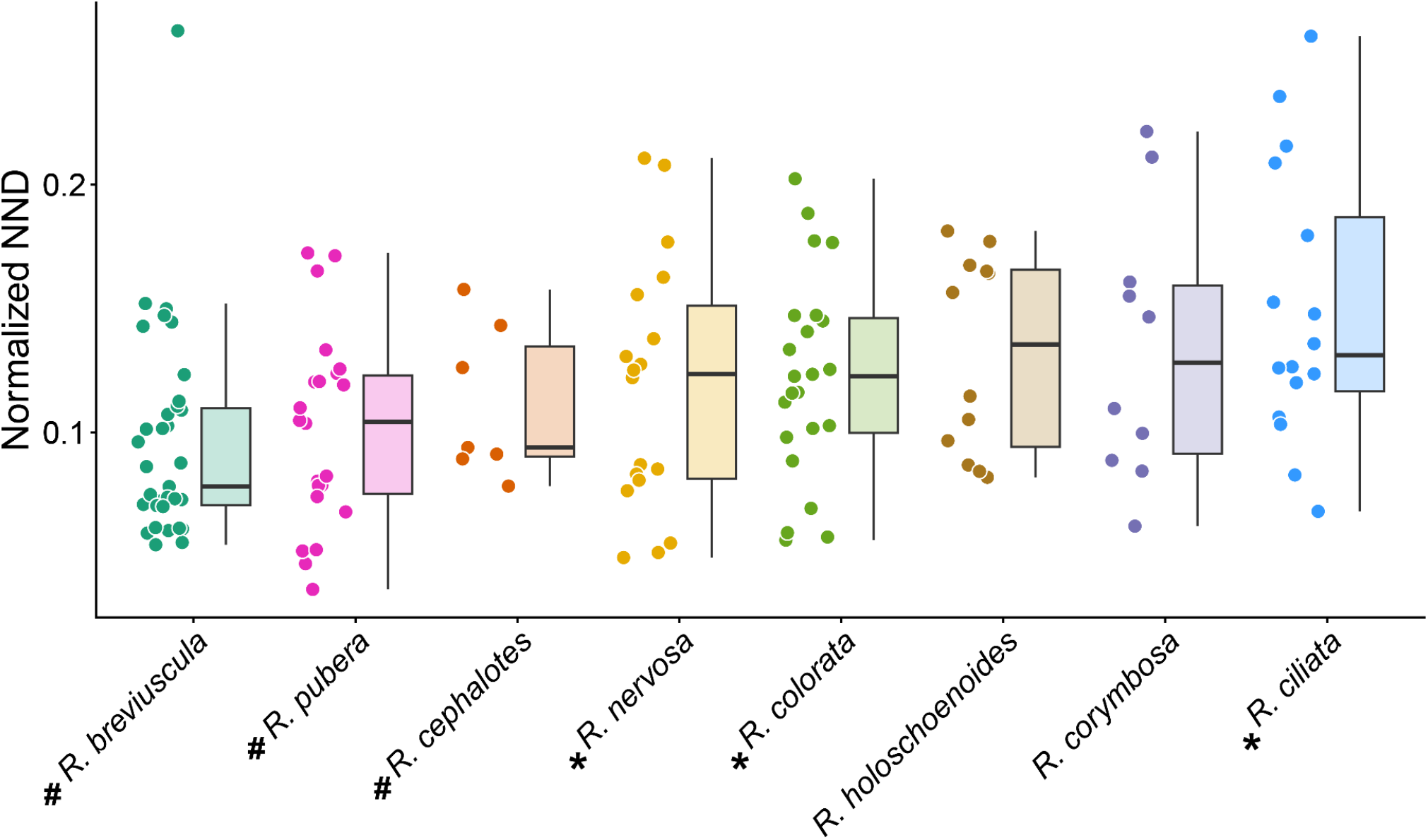
Quantification of nearest-neighbour distances (NND) between telomere FISH signals across species. Values are given as the average NND normalised by the square root of cell area. Boxplots show median, interquartile range, and distribution of NND values. Species with distal-biased CO landscapes (#) showed the smallest NNDs compared with species showing irregular recombination patterns (*).

**Supplementary Fig. 8:**
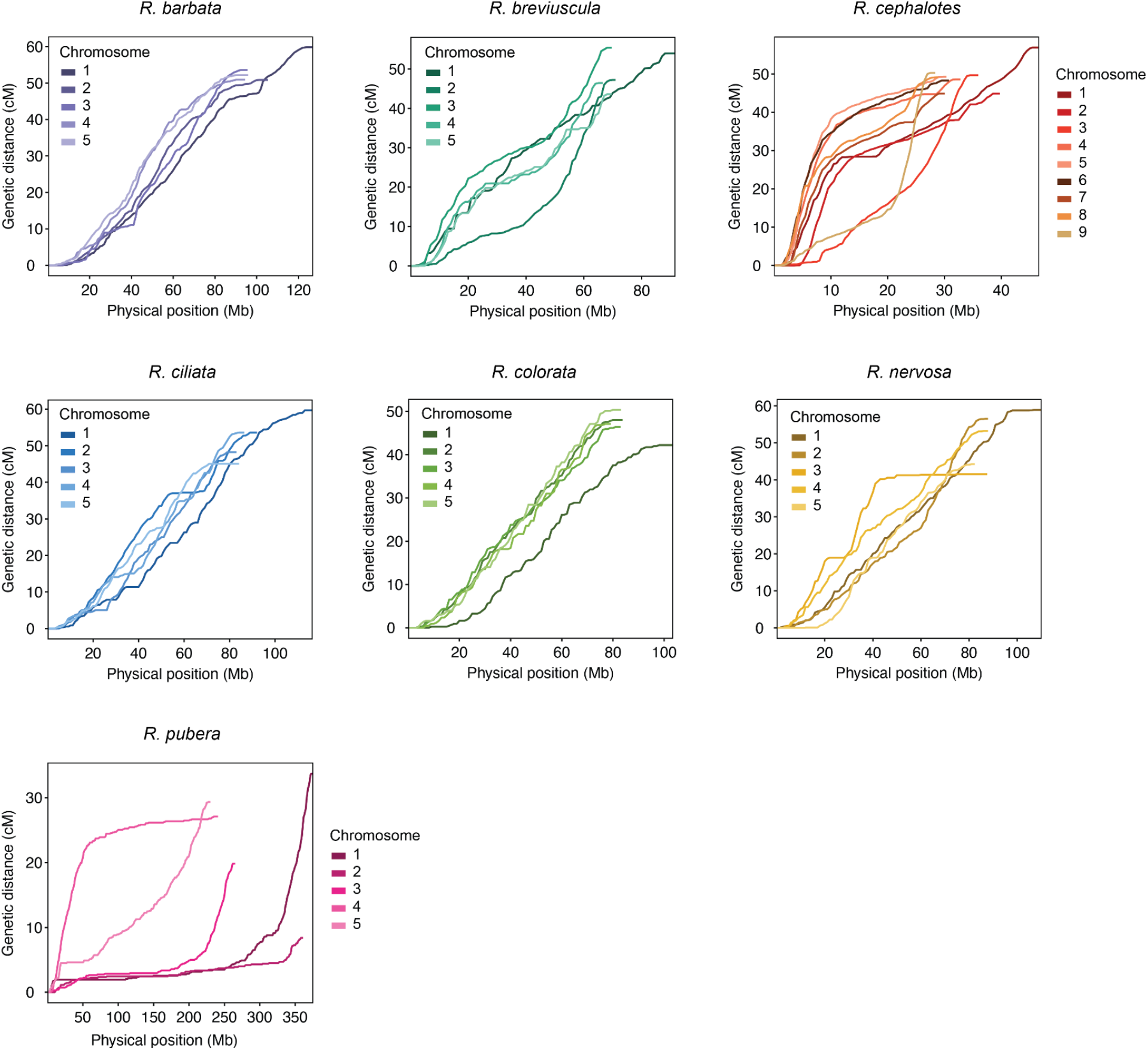
Marey maps distinguish distal-biased and irregular recombination landscapes across *Rhynchospora*. Marey maps showing cumulative genetic distance (cM) as a function of physical position (Mb) for all chromosomes in seven *Rhynchospora* species (*R. barbata, R. breviuscula, R. cephalotes, R. ciliata, R. colorata, R. nervosa*, and *R. pubera*). Individual curves represent chromosomes within each species. Species with distal-biased recombination landscapes display steep increases in genetic distance toward one or both chromosome ends, whereas species with irregular landscapes show more heterogeneous, segmental slopes along chromosome arms. Across both landscape classes, smaller chromosomes consistently exhibit higher recombination density than larger chromosomes, highlighting chromosome size as a quantitative determinant of recombination despite qualitative differences in CO distribution.

**Supplementary Fig. 9:**
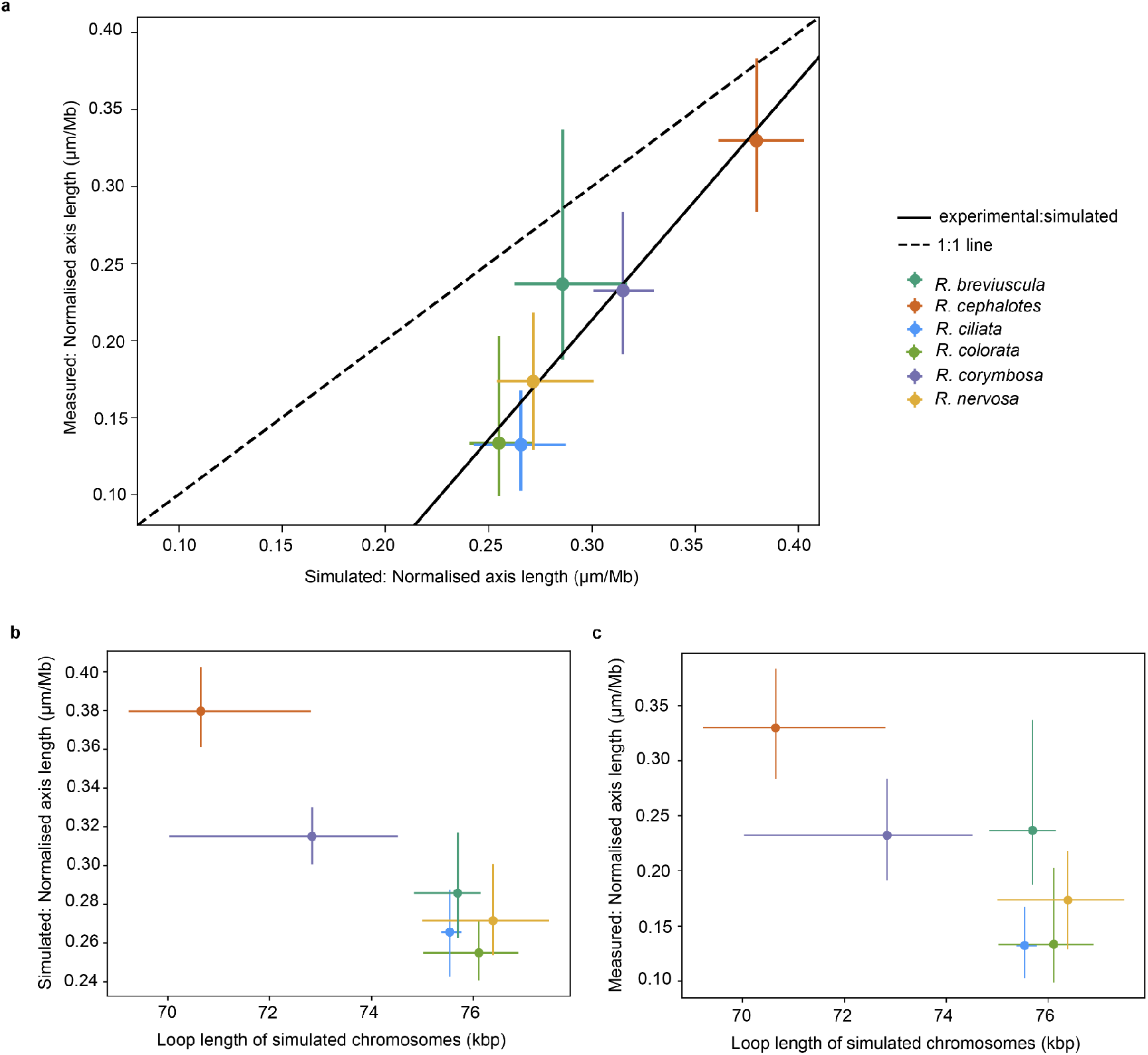
Chromosome loop length and karyotype configuration predict meiotic axis length across *Rhynchospora*. (**a**) Comparison between experimentally measured normalised chromosome axis length (µm/Mb) and simulated axis length derived from species-specific karyotype configurations across six *Rhynchospora* species. Points represent species means, with horizontal and vertical error bars indicating measured samples and simulated replicas variability. The solid line shows the linear fit between measured and simulated values, while the dashed line indicates a 1:1 relationship, demonstrating close correspondence between empirical measurements and model predictions. (**b**) Relationship between loop length of simulated chromosomes and simulated normalised axis length, showing that shorter loops are associated with longer axis length per megabase. (**c**) Relationship between loop length of simulated chromosomes and experimentally measured normalised axis length, confirming that loop size quantitatively predicts meiotic axis length in vivo.

## Supplementary Datasets

**Supplementary Dataset 1**. Hi-C contact matrices, BUSCO assessment and *k*-mer estimation of genome size and heterozygosity for all *Rhynchospora* species included in the pangenome.

**Supplementary Dataset 2**. List of identified chromosome fusion and fission junctions across the *Rhynchospora* pangenome, including breakpoint coordinates and repeat associations.

**Supplementary Dataset 3**. Genome-wide crossover positions, CO frequencies, and associated genomic feature densities for all species with recombination maps.

**Supplementary Dataset 4**. Hi-C–derived chromatin contact probability profiles and inferred loop/domain size estimates used for axis-length modelling.

